# The Female Side of Autism: Sexual Dichotomies impacting the Helsmoortel-Van der Aa syndrome pathology

**DOI:** 10.64898/2026.09.20.752986

**Authors:** Claudio Peter D’Incal, Joe Ibrahim, Mathijs B. Van der Lei, Lusine Harutyunyan, Kim Van Meel, Ellen Elinck, Ayu Scott, Anne Schepers, Lauren Moons, Marie Hannaert, Zlatko Marusic, Mirna Anicic, Jurica Vukovic, Paschalis Theotokis, Nikolaos Grigoriadis, Monique de Booij Oomen, Ligia Mateiu, Erik Fransen, Anna C Jansen, Marije Meuwissen, Dale John Annear, Illana Gozes, R. Frank Kooy

## Abstract

**Background:** Helsmoortel-Van der Aa syndrome, caused by pathogenic variants in *ADNP*, is characterised by substantial clinical heterogeneity, but how biological sex influences disease biology and treatment response remains an emerging area of investigation. Here, we investigated whether sex modifies the behavioural and molecular consequences of *Adnp* deficiency, the response to the investigational drug candidate davunetide (NAP), and the clinical phenotype of Helsmoortel-Van der Aa syndrome.

**Methods:** We studied male and female *Adnp* heterozygous mice harbouring the p.Leu822Hisfs*6 variant and sex-matched wild-type littermates using continuous 24-h behavioural phenotyping, hippocampal genome-wide DNA methylation and bulk RNA sequencing. NAP effects on behaviour, hippocampal ADNP protein abundance, and epi-transcriptomic responses were assessed using genotype-by-treatment models. Four Core Genotypes mice were assessed to distinguish sex-chromosome and gonadal contributions to hippocampal *Adnp* transcript expression, and reproductive-tissue transcriptomes were analysed in *Adnp* mice to unravel a possible endocrine component of the disease. Findings were compared with clinical and developmental data from 129 individuals with Helsmoortel-Van der Aa syndrome.

**Findings:** *Adnp* deficiency produced a shared overall behavioural phenotype in male and female mice, but through distinct patterns of behavioural disruption, indicating that sex modifies the implementation rather than the magnitude of the phenotype. This distinction was also evident at the molecular level: hippocampal DNA methylation was predominantly determined by genotype and showed convergence between sexes, whereas transcriptional responses diverged, with mitochondrial pathways predominating in males and chromatin, RNA processing, and cell-cycle pathways in females. NAP partially shifted behavioural abnormalities towards the wild-type state in both sexes and increased hippocampal ADNP protein abundance in heterozygous mice, without reversing the underlying methylation phenotype. Its transcriptional effects were markedly broader in males, indicating a sex-dependent molecular response to treatment. In reproductive tissues, *Adnp* deficiency was associated with convergent alterations in steroid hormone biosynthesis, despite no evidence for direct regulation of hippocampal *Adnp* expression by sex-chromosome complement or gonadal hormonal state. By contrast, sex was not associated with robust differences across the clinical and developmental features assessed in 129 individuals with Helsmoortel-Van der Aa syndrome after correction for multiple testing.

**Interpretation:** Biological sex modifies how *Adnp* deficiency is expressed at behavioural and molecular levels, but these differences do not necessarily define distinct clinical phenotypes. The convergence of the methylation response alongside greater divergence in transcriptional and behavioural responses identifies sex as a modifier of disease biology rather than a determinant of the core phenotype. The sex-dependent response to NAP further indicates that biological sex may influence pharmacological responses even when treatment-associated behavioural improvement is observed in both sexes. These findings support consideration of biological sex as a prespecified variable in preclinical and clinical studies of Helsmoortel-Van der Aa syndrome and suggest that systematic investigation of sex-dependent biology may also be relevant to therapeutic development across rare diseases.

## Introduction

Neurodevelopmental disorders (NDDs) comprise a heterogeneous group of genetically complex conditions, with reported prevalence increasing alongside improvements in paediatric screening and diagnosis (1). Developmental delay (DD), intellectual disability (ID), autism spectrum disorder (ASD), attention-deficit hyperactivity disorder (ADHD), and motor developmental disorders exhibit substantial clinical and genetic heterogeneity and show pronounced sex-dependent differences in prevalence, symptom presentation, and disease trajectories (2). Increasing evidence indicates that biological sex can influence disease penetrance, phenotypic expressivity, and genetic burden across NDDs, including rare monogenic conditions, although these effects remain insufficiently characterised in ultra-rare ASD-ID syndromes (3,4). In contrast, large-scale exome sequencing studies have generally reported a near-equal sex distribution among individuals carrying pathogenic variants in highly penetrant NDD genes, with only a limited number of genes showing evidence of male- or female-biased enrichment (5).

*De novo* variants in the *Activity-Dependent Neuroprotective Protein (ADNP)* gene cause Helsmoortel-Van der Aa syndrome (OMIM, 615873), a rare ASD-ID disorder characterised by multisystemic involvement (6). ADNP functions as a chromatin-associated transcription factor (7) and is essential for neocortical neurogenesis (8). The protein contains nine zinc finger motifs, the neuroprotective octapeptide sequence <u>NAP</u>VSIPQ (the active peptide of the investigational drug davunetide), a bipartite nuclear localisation signal (NLS), and a homeobox domain, collectively supporting its predominant nuclear regulatory function (9). In addition to ASD and ID, individuals with Helsmoortel-Van der Aa syndrome typically present with facial dysmorphisms, cardiac abnormalities, behavioural disturbances, feeding and gastrointestinal problems, visual impairments, and additional multisystem health complications (10,11). Clinical characterisation of a Belgian patient cohort demonstrated an approximately equal sex distribution, consistent with large-scale exome sequencing studies indicating no apparent sex bias among individuals with pathogenic *ADNP* variants (5,10). The prominent genetic contribution of *ADNP* to NDDs was further supported by a large-scale analysis of *de novo* variants across 11 ASD or DD cohorts, comprising 46,612 parent-proband trios, together with 6,557 ASD trios from the Simons Foundation Powering Autism Research for Knowledge (SPARK) cohort. Across three independent statistical models, *ADNP* showed the strongest enrichment for *de novo* variants, alongside *ARID1B*, *DYRK1A*, *POGZ*, and *CTNNB1* (5). Despite the increasing recognition of sex-dependent effects across NDDs, sex-stratified phenotypic analyses in Helsmoortel-Van der Aa syndrome remain limited (9–11).

Preclinical studies provide evidence that biological sex may influence the phenotypic consequences of *Adnp* deficiency, although findings differ between models. Sex-dependent phenotypes have been reported in haploinsufficient *Adnp* mice (Δexon 1-6) (12–14) and in mice harbouring the p.Tyr718* nonsense *Adnp* variant affecting the NLS domain (15,16). In contrast, an independent haploinsufficient *Adnp* mouse model (Δexon 6) did not show pronounced sex-dependent effects. This discrepancy may reflect differences in the genomic extent of *Adnp* disruption between the models, as well as differences in the age at which phenotypes were assessed (17). Beyond loss-of-function models, sex-dependent regulation of *Adnp* expression has been reported in the murine hypothalamic arcuate nucleus, with females showing higher immunoreactivity than males in an oestrous-cycle-dependent manner (18). Homozygous *Adnp* deficiency results in embryonic lethality at E8.5-9 (19), while transcriptomic microarray analyses of homozygous embryos implicated *Adnp* in pathways associated with steroid hormone biosynthesis and endocrine regulation (20).

Recently, we generated a novel heterozygous (HET) *Adnp* mouse model harbouring the p.Leu822Hisfs*6 frameshift variant located within a mutational hotspot for pathogenic human *ADNP* variants. Behavioural and molecular analyses of 10-week-old male mice using the Live Mouse Tracker (LMT) platform identified autism-associated phenotypes together with dysregulation of Wnt signalling and cytoskeletal pathways implicated in Helsmoortel-Van der Aa syndrome pathology (21). The LMT platform enables continuous, automated behavioural phenotyping in socially housed mice and provides a framework for longitudinal assessment of behavioural responses to therapeutic interventions (22). Importantly, the ADNP-derived investigational drug davunetide (NAP, also known as CP201) has shown preclinical efficacy across multiple *Adnp*-deficient models (23). This provides an opportunity to determine whether therapeutic responsiveness to NAP differs according to biological sex in our *Adnp* frameshift model (21,22).

Using this *Adnp* frameshift mouse model, we aimed to define sex-dependent behavioural and molecular alterations in the hippocampus and determine whether these differences influence therapeutic responsiveness. We investigated sex differences in continuous 24-hour behavioural profiles, hippocampal epigenomic and transcriptomic signatures, and gonadal molecular pathways, with translational validation in human gonadal tissues and clinical phenotyping in a large Helsmoortel-Van der Aa syndrome cohort. We further assessed whether the investigational drug davunetide (NAP) exerts sex-dependent therapeutic effects. By integrating behavioural, molecular, endocrine-associated, translational, and clinical analyses across murine and human systems, this study investigates biological sex as a potential modifier of disease mechanisms and therapeutic response in Helsmoortel-Van der Aa syndrome.

## Materials and methods

### Animals

Male and female *C57BL/6JCr-Adnp^em1Ant^/J* HET mice and their wild-type (WT) littermates (The Jackson Laboratory; RRID:IMSR_JAX:033128) were generated by the Cognitive Genetics laboratory, Centre of Medical Genetics, Antwerp University Hospital (UZA)/University of Antwerp (UA). Animals were socially housed in standard mouse cages (22.5 cm x 16.7 cm x 14 cm) with a maximum of eight animals per cage under controlled temperature and humidity conditions and a 12 h light-dark cycle, with food and water available *ad libitum*. Environmental enrichment consisted of a platform, tunnel and cotton nesting material. All experiments were performed in 10-week-old mice. All animal procedures complied with European Directive 2010/63/EU and were approved by the Animal Ethics Committee of the University of Antwerp (ECD 2023-17). Experiments were conducted in accordance with the ARRIVE guidelines. Genotyping was performed as described previously (21).

### NAP treatment

NAP, formulated as davunetide-acetate, was custom synthesised and provided under material transfer agreement by Ramot at Tel Aviv University (Israel). Treatment allocation was performed at the litter level immediately after birth, before determination of offspring sex and genotype. Entire litters were randomly assigned to receive either NAP or DD-vehicle treatment (14,15), and allocation was maintained throughout the experimental period. Genotype and sex were determined at weaning (postnatal day P28), after which animals were stratified into the experimental groups. During early postnatal development (postnatal days P0-P21), pups received daily subcutaneous injections of either vehicle solution (phosphate-buffered saline, PBS) or NAP (25 µg/mL dissolved in PBS). Injection volumes were adjusted according to body weight as follows: 20 µL (P0-4), 40 µL (P5-10), 80 µL (P11-14), 120 µL (P15-18), and 160 µL (P19-21). Following weaning, mice received daily intranasal administration of either NAP or DD-vehicle solution (5 µL per nostril). The DD-vehicle solution consisted of 129 mM NaCl, 8 mM citric acid monohydrate, 17 mM disodium phosphate dihydrate and 0.01 % benzalkonium chloride. Intranasal administration was performed in manually restrained mice using a P10 pipette fitted with sterile filter tips (24). Animals were monitored daily throughout the treatment for general health and welfare. Body weight was recorded longitudinally following NAP or DD-vehicle administration. Behavioural and molecular analyses were performed using predefined analytical pipelines, and investigators were blinded to genotype during downstream data acquisition and quantitative analyses where feasible.

### Sample size and experimental design

Sample sizes for the individual experimental procedures are summarised in table 1. Where applicable, group sizes were determined *a priori* using G∗Power (version 3.1) to achieve 80% statistical power at α = 0.05, based on effect sizes reported in previous studies of *Adnp*-deficient mouse models, behavioural phenotyping, and multi-omics analyses. All animal experiments were designed in accordance with the principles of reduction, refinement and replacement. For transcriptomic and epigenomic analyses, sample sizes were selected in accordance with established experimental designs for controlled murine bulk RNA sequencing and DNA methylation studies. Human reproductive-tissue analyses were exploratory and constrained by the rarity of Helsmoortel-Van der Aa syndrome and the limited availability of patient-derived tissues. All available patient-derived specimens and matched control tissues were included. The clinical cohort analysis used all individuals for whom sex-stratified data were available from the previously published cohort (11).

**Table 1.** Sample size and justification for each experimental procedure. Sample sizes for animal experiments were determined according to the study design and, where applicable, informed by prior studies and power calculations. Animal experiments were conducted in accordance with the principles of reduction, refinement and replacement (3Rs).

| Experiment | Sample size | Justification for sample size | References |
| --- | --- | --- | --- |
| *NAP treatment and body-weight determination* | **Males:**<br>WT + DD (n = 14)<br>HET + DD (n = 12)<br>WT + NAP (n = 12)<br>HET + NAP (n = 11)<br>**Females:**<br>WT + DD (n = 11)<br>HET + DD (n = 11)<br>WT + NAP (n = 13)<br>HET + NAP (n = 11) | Group sizes were selected based on previous behavioural and therapeutic studies in *Adnp*-deficient and related neurodevelopmental mouse models. Group sizes of approximately 10-15 animals per condition were considered appropriate to detect moderate genotype-, treatment- and sex-associated effects while accounting for variability introduced by sex-stratified analyses. | (14,15,24–27) |
| *Live Mouse Tracker behavioural phenotyping* | **Males:**<br>WT + DD (n = 14)<br>HET + DD (n = 12)<br>WT + NAP (n = 12)<br>HET + NAP (n = 11)<br>**Females:**<br>WT + DD (n = 11)<br>HET + DD (n = 11)<br>WT + NAP (n = 13)<br>HET + NAP (n = 11) | The same treatment cohorts were used for continuous automated behavioural phenotyping. Group sizes were selected to permit genotype-, treatment- and sex-stratified analyses while providing adequate sensitivity to detect moderate behavioural effects. | (21,22) |
| *Hippocampal Immunoblotting* | **Males:**<br>WT + DD (n = 6)<br>HET + DD (n = 6)<br>WT + NAP (n = 6)<br>HET + NAP (n = 6)<br>**Females:**<br>WT + DD (n = 6)<br>HET + DD (n = 6)<br>WT + NAP (n = 6)<br>HET + NAP (n = 6) | Group sizes were informed by previous biochemical studies investigating NAP-related molecular effects in mouse models and were selected to detect moderate-to-large genotype- and treatment-associated changes in protein abundance. | (24,26) |
| *Hippocampal RT-qPCR expression analysis* | **Males:**<br>WT + DD (n = 7)<br>HET + DD (n = 7)<br>WT + NAP (n = 7)<br>HET + NAP (n = 7)<br>**Females:**<br>WT + DD (n = 7)<br>HET + DD (n = 7)<br>WT + NAP (n = 7)<br>HET + NAP (n = 7) | RT-qPCR was performed to assess *Adnp* transcript abundance and selected sex- and treatment-associated transcripts and to provide targeted validation of representative findings from bulk RNA sequencing. | (21,28) |
| *Hippocampal Infinium Mouse Methylation BeadChip analysis* | **Males:**<br>WT + DD (n = 7)<br>HET + DD (n = 7) | Sample sizes were determined *a priori* using G*Power (80% power, $\alpha = 0.05$) and were consistent with previous methylome-profiling studies in | (29,30) |
|  | WT + NAP (n = 7)<br>HET + NAP (n = 7)<br>**Females:**<br>WT + DD (n = 7)<br>HET + DD (n = 7)<br>WT + NAP (n = 7)<br>HET + NAP (n = 7) | neurodevelopmental mouse models. Group sizes were selected to detect biologically relevant methylation differences across genotype, treatment and sex. |  |
| Targeted pyrosequencing validation | Same animals as hippocampal Infinium Mouse Methylation BeadChip array | Targeted pyrosequencing was performed as an orthogonal validation of selected differentially methylated loci identified by array-based methylation profiling. | (29,30) |
| Bulk hippocampal RNA sequencing | **Males:**<br>WT + DD (n = 5)<br>HET + DD (n = 4)<br>WT + NAP (n = 5)<br>HET + NAP (n = 4)<br>**Females:**<br>WT + DD (n = 6)<br>HET + DD (n = 5)<br>WT + NAP (n = 5)<br>HET + NAP (n = 5) | Bulk RNA sequencing was performed to characterise genotype-associated transcriptional alterations separately in males and females and to assess treatment-associated transcriptional effects. Sample sizes were informed by previous bulk transcriptomic studies in neurodevelopmental mouse models and were selected to provide adequate sensitivity for differential-expression analyses at an FDR threshold of 0.05. | (14,16,21,31) |
| Reproductive-tissue bulk RNA sequencing | **Testis:**<br>WT (n = 4)<br>HET (n = 4)<br>**Uterus:**<br>WT (n = 4)<br>HET (n = 4) | Exploratory bulk RNA sequencing was performed to determine whether *Adnp* deficiency was associated with transcriptional alterations in reproductive tissues and to identify pathways that might represent a shared endocrine or reproductive component of the molecular phenotype. Four biological replicates per genotype were analysed for each tissue. | (31) |
| Reproductive-tissue RT-qPCR validation | Same biological samples as used for reproductive-tissue bulk RNA sequencing. | Targeted RT-qPCR was performed on the same biological samples to validate representative genotype-associated transcripts identified by bulk RNA sequencing, including genes selected for their relevance to tissue differentiation and reproductive function. | (28) |
| Publicly available hippocampal RNA sequencing: Four Core Genotypes model (GSE184098) | **Testis:**<br>XY-Sry (n = 6)<br>XXSry (n = 6)<br>**Ovaries:**<br>XX (n = 6)<br>XY- (n = 5) | Publicly available hippocampal RNA-sequencing data from the Four Core Genotypes model at the age of 12 months were analysed to distinguish effects associated with sex chromosome complement from those associated with gonadal hormonal state and to assess whether *Adnp* expression was directly or indirectly associated with sex- or hormone-dependent transcriptional programmes. | (32) |
| Human reproductive-tissue RT-qPCR, immunohistochemistry and immunoblotting | **Testis:**<br>Control subjects (n = 3)<br>HVDAS (n = 1)<br>**Uterus:**<br>Control subjects (n = 3)<br>HVDAS (n = 1) | Sample availability determined the cohort size. Given the rarity of HVDAS and the limited availability of human reproductive tissue, all available patient specimens were included. Three independent control specimens per tissue were analysed to provide an exploratory comparison with HVDAS tissue. | (33–35) |
| *Clinical cohort re-analysis* | **HVDAS** (n = 129; 69 males and 60 females) | Clinical analyses were performed using all available phenotypic and demographic data from the largest currently available Helmsmoortel-Van der Aa syndrome cohort to assess sex-dependent clinical manifestations and translational relevance of preclinical findings. | (10,11) |

### Live mouse tracker

Live Mouse Tracker (LMT) recordings were performed as previously described (21). Briefly, the LMT system combines infrared depth imaging, machine learning-based behavioural classification and radio-frequency identification (RFID)-based animal identification to enable continuous tracking and automated behavioural phenotyping of group-housed mice. Mice were recorded continuously for 24 h under controlled environmental conditions across independent recording sessions. Behavioural events were automatically extracted using the LMT framework and analysed using the MouseKing behavioural analysis pipeline (https://github.com/DaleAnnear/MouseKing.git). Thirty-three individual and social behavioural features were quantified, including exploratory behaviour, locomotion, rearing, stretched-attend posture (SAP), social contacts, group formation, following behaviour, social approaches and oral-oral and oral-genital interaction sequences. Behavioural analyses were performed in sequential stages. First, unsupervised multivariate analysis was performed across male and female WT and HET mice receiving DD-vehicle to identify genotype- and sex-dependent behavioural phenotypes. Baseline behavioural phenotypes were subsequently analysed separately in males and females. Finally, NAP responsiveness was assessed within the sex-stratified experimental groups. Behavioural events shorter than 15 frames were excluded during quality control. Data were normalised using z-score transformation and within-cage averaging to minimise batch and housing effects. Principal component analysis was used to assess multivariate behavioural structure. Behavioural variables with hierarchical structure were analysed using linear mixed-effects models, with genotype and treatment included as fixed effects and cage identity as a random intercept. The 33 LMT behavioural variables were assigned to one of the five predefined behavioural domains. Absolute genotype-associated effect sizes (Cohen’s *d*) were summed within each domain and expressed as the proportion of the summed absolute effect size across all behavioural variables.

### Immunoblotting

Hippocampal, testicular and uterine tissues were dissected on ice, snap frozen in liquid nitrogen and stored at −80°C until processing. Tissue samples were homogenised using a TissueRuptor II homogeniser (Qiagen, Germany) in ice-cold radioimmunoprecipitation assay (RIPA) buffer containing 150 mM NaCl, 50 mM Tris-HCl, 0.5% sodium deoxycholate, 1% NP-40, and 2% SDS, supplemented with protease inhibitor cocktail (cOmplete Mini EDTA-free, Roche; 04693159001) and PhosSTOP phosphatase inhibitor (Roche; 4906845001). Lysates were incubated for 1 h at 4°C under agitation and clarified by centrifugation at maximum speed for 30 min at 4°C. Protein concentrations were determined using the Pierce BCA Protein Assay Kit (Thermo Fisher Scientific; 23225). Immunoblotting was performed as previously described (21,36). Briefly, 20 µg total protein per sample was reduced, denatured for 10 min at 70°C, separated on Bolt 4-12% Bis-Tris polyacrylamide gels (Invitrogen; NW04120BOX) and transferred onto nitrocellulose membranes using wet electrotransfer. Membranes were blocked and incubated with primary antibody against ADNP (Abcam; RRID:AB_3097704), CNN1 (Abcam; RRID:AB 2291941), and TSPAN8 (Abcam; RRID:AB_3751514), followed by species-appropriate horseradish peroxidase (HRP)-conjugated secondary antibodies (Agilent Technologies, USA). Protein bands were visualised using SuperSignal West Femto Maximum Sensitivity Substrate (Thermo Fisher Scientific; 34095) on an Amersham Imager 800 system (Cytiva, USA). Densitometric quantification was performed using ImageJ. Statistical analyses were conducted in GraphPad Prism (version 10.6.1) using two-way ANOVA followed by Tukey’s multiple comparisons test or Welch’s t-test. Full uncropped immunoblots are provided in additional file S1.

### Quantitative real-time PCR

Total RNA was extracted from hippocampal tissue of 10-week-old male and female WT and HET mice receiving NAP or DD-vehicle, as well as from murine and human testicular and uterine tissues, using the RNeasy Mini Kit (Qiagen; 74106) according to the manufacturer’s instructions. RNA concentration was quantified using the Qubit RNA Broad Range Assay Kit (Invitrogen; Q10211). One microgram of RNA was reverse transcribed into complementary DNA (cDNA) using the SuperScript III Reverse Transcriptase Kit (Invitrogen; 18080093) according to the manufacturer’s instructions. Primer efficiencies were assessed using standard dilution curves generated from pooled WT and HET cDNA samples. Primer pairs with amplification efficiencies between 90% and 110% were retained for analysis. Quantitative real-time polymerase chain reaction (RT-qPCR) was performed in technical triplicate using Takyon No ROX SYBR 2× MasterMix (Eurogentec; UF-NSMT-B0701) on the CFX384 Touch Real-Time PCR Detection System (Bio-Rad, USA). Primer sequences are provided in additional file S2. Relative gene expression was quantified using qbase+ software (Biogazelle, Belgium), applying a maximum allowable replicate deviation of 0.5 cycle threshold units between technical triplicates.

For validation of hippocampal RNA-sequencing findings, representative genes were selected based on robust genotype-associated differential expression and biological relevance to the transcriptional pathways identified by genome-wide analysis. Hippocampal validation data were analysed using Welch’s t-test or two-way ANOVA followed by Šídák’s multiple-comparisons test.

For validation of reproductive-tissue RNA-sequencing findings, representative transcripts were selected to capture genotype-associated alterations in genes with established relevance to reproductive tissue function, cellular differentiation and tissue-specific biology. Adh1 and Tspan8 were selected for testicular validation, whereas Cnn1 and Tgfbr3l were selected for uterine validation. Gonadal comparisons between WT and HET mice were performed using Welch’s t-test.

ADNP transcript abundance was further assessed using complementary transcript-specific assays designed to distinguish total Adnp transcript abundance from transcripts retaining the wild-type sequence and transcripts spanning the deleted region. These assays were performed as previously described (21).

### Infinium mouse methylation BeadChip array and data processing

Total DNA was isolated from the hippocampal tissue of male and female WT and HET mice receiving NAP or DD-vehicle using the DNeasy Blood and Tissue kit (Qiagen; 69504) according to the manufacturer’s instructions. Bisulphite conversion of 200 ng genomic DNA was performed using the EZ DNA Methylation Kit (Zymo Research; D5001). Successful conversion was verified by amplification of a methylation-conserved fragment of the murine *Line1* gene using the PyroMark PCR kit (Qiagen; 978703) with the following primers: forward 5′-GCGCGAGTCGAAGTAGGGC-3′ and reverse 5‘-ACCCAACGATACCTAATAATAAAACC-3’. PCR products were verified by tris-boric acid-EDTA (TBE) electrophoresis on 1.5% agarose gels.

Bisulphite-converted DNA was hybridised to the Infinium Mouse Methylation EPIC BeadChip (Illumina; 20041558) according to the manufacturer’s protocol and scanned using the Illumina Hi-Scan system. CpG annotation and genomic context relative to CpG islands, shores and shelves were obtained from the Illumina BeadChip manifest and based on *Mus musculus* genome build GRCm38.p6/mm10. Raw IDAT files were processed in *R* (version 4.3.1) using the Bioconductor package ENmix (version 1.42.0). Signal intensities were corrected for background and dye bias using out-of-band background correction and RELIC normalisation, respectively, followed by quantile normalization and probe-type bias adjustment using regression on correlated probes. Cross-reactive probes, probes containing single nucleotide polymorphisms (SNPs) and probes mapping to the sex chromosomes were excluded. β-values were subsequently calculated for all retained probes. Differential methylation was assessed using the non-parametric Wilcoxon rank-sum test. To prioritise biologically meaningful methylation changes, CpG probes were considered biologically differentially methylated when the absolute methylation difference was at least 10% (|Δβ| ≥ 0.1), corresponding to hypomethylation (Δβ-values < -0.1) and hypermethylation (Δβ-values > 0.1) (37).

Methylation analyses were performed separately in males and females to define sex-stratified genotype-associated methylation signatures. Baseline genotype-associated methylation was assessed by comparing DD-vehicle-treated WT and HET mice within each sex. Genes associated with CpG probes meeting the predefined effect-size threshold were annotated and used for downstream functional analyses. To assess convergence between male and female genotype-associated methylation signatures, unique genes associated with differentially methylated probes were compared between sexes. The number and proportion of shared genes were calculated, together with the Jaccard index. For statistical assessment of overlap, the gene universe was restricted to genes tested in both sexes, and Fisher’s exact test was used to determine whether the observed overlap exceeded that expected by chance. To investigate the effect of NAP treatment on hippocampal DNA methylation, NAP-treated animals were compared with DD-vehicle-treated animals within genotype- and sex-matched groups. Linear models including genotype, treatment and the genotype × treatment interaction were then fitted to identify treatment-associated methylation changes while accounting for genotype-dependent effects. The interaction term was evaluated first to determine whether the effect of NAP differed according to genotype. Where the interaction term was not significant, reduced models including genotype and treatment as main effects were fitted to evaluate treatment-associated methylation changes independent of genotype. CpG probes showing significant genotype × treatment interactions were considered separately and were not interpreted as a uniform treatment effect. Multiple testing was controlled using the Benjamini-Hochberg false discovery rate (FDR) procedure. Genes associated with CpG probes meeting the |Δβ| ≥0.1 threshold were used for functional interpretation. Gene ontology and pathway enrichment analyses were performed using Metascape with default parameters and multiple-testing correction (38). Enrichment terms with adjusted p < 0.05 were considered statistically significant.

### Targeted pyrosequencing analysis

Targeted pyrosequencing was performed to validate biologically relevant differentially methylated loci identified in the genome-wide hippocampal methylation analysis. Candidate loci were prioritised according to predefined criteria, including an absolute methylation difference of at least 10%, localisation within annotated genes or regulatory regions, biological relevance to pathways identified by genome-wide analyses, and representation of shared or sex-associated methylation changes. Forward, reverse and sequencing primers were designed using PyroMark Assay Design 2.0 software (Qiagen, Germany) (additional file S3). Bisulphite-converted DNA was PCR amplified using the PyroMark PCR kit (Qiagen, Germany), and amplification was confirmed by agarose gel electrophoresis. Biotinylated PCR products were immobilised on streptavidin-coated Sepharose beads (GE Healthcare, USA), purified and denatured using the PyroMark Q24 Vacuum Workstation, and annealed to sequencing primers for 5 min at 80°C. Pyrosequencing was performed using the PyroMark Q24 instrument (Qiagen, Germany), and methylation percentages were quantified using PyroMark Q24 software. Statistical analyses were performed in GraphPad Prism (version 10.6.1).

### Bulk transcriptome sequencing

Bulk RNA sequencing was performed on hippocampal tissue and on mouse testis and uterus to characterise tissue-specific transcriptional consequences of *Adnp* deficiency and, in hippocampal samples, to assess genotype- and treatment-associated effects. Samples with an RNA integrity number (RIN) > 6.5 were selected for sequencing and submitted to Novogene (Munich, Germany). Libraries were prepared using the Novogene NGS RNA Library Prep Set (PT042) and sequenced on an Illumina NovaSeq 6000 platform using a paired-end 150 bp (PE150) strategy. Raw sequencing reads were subjected to quality control and adapter trimming before alignment to the *Mus musculus* reference genome (GRCm38.p6, Ensembl version 102) using STAR. Gene-level read counts were generated using featureCounts from the Subread package. Differential gene expression analyses were performed in *R* using DESeq2. For hippocampal analyses, genotype-associated transcriptional changes were assessed separately in males and females by comparing WT and HET littermates receiving DD-vehicle. These genotype comparisons were evaluated using the DESeq2 Wald test. To assess whether the transcriptional response to NAP treatment differed by genotype, a genotype × treatment interaction term was included in the model and evaluated using the DESeq2 likelihood ratio test (LRT). Genes with an FDR-adjusted p < 0.05 were considered differentially expressed. Hierarchical clustering was performed using differentially expressed genes to assess genotype-associated transcriptional structure. Functional enrichment analyses were performed using gene set enrichment analysis implemented in *R* using GSEA, focusing on Gene Ontology biological processes and curated pathway databases. To quantify convergence between male and female genotype-associated transcriptional responses, the DEG sets were compared, and the number and proportion of shared genes and the Jaccard index were calculated. Statistical enrichment of the observed overlap was assessed using Fisher’s exact test after restricting the background universe to genes tested in both sexes. This analysis used the common gene universe of 16,508 genes.

### Four Core Genotypes analysis

To investigate whether *Adnp* expression was influenced by sex chromosome or gonadal hormonal state independently of the *Adnp* genotype, publicly available hippocampal RNA-sequencing data from the Four Core Genotypes (4CG) mouse model at the age of 12 months were analysed (GEO accession: GSE184098). The 4CG model separates chromosomal sex from gonadal sex through translocation of the *Sry* gene. XX and XY mice lacking *Sry* develop ovaries, whereas XX and XY mice carrying the *Sry* transgene develop testes. This design permits assessment of the independent contributions of sex chromosome complement and gonadal sex to hippocampal gene expression (32). Normalised *Adnp* expression was analysed using linear mixed-effects models incorporating sex chromosome complement and gonadal sex as fixed effects, with cage as a random effect. Multiple testing was controlled using FDR correction. To assess whether *Adnp* might indirectly contribute to sex- or hormone-associated transcriptional programmes through its reported transcription-factor activity (35), DEGs associated with sex- and hormonal-state differences in the 4CG hippocampus were subjected to transcription-factor enrichment analysis using ChEA3. The overlap between the identified DEG set and genes associated with the *Adnp* regulon was assessed, and the relative enrichment rank of *Adnp* among queried transcription factors was recorded.

### Hormone-associated hippocampal gene expression

To assess whether the female hippocampal cohort showed evidence of a pronounced high-oestrogen transcriptional state at the time of tissue collection, expression of *Esr1*, *Pgr* and *Bdnf* was examined in the same bulk RNA-sequencing cohort. Male mice were used as a non-cycling reference. Normalised gene counts were compared across sex, genotype and treatment groups using linear mixed-effects models, with FDR correction for multiple comparisons.

### Human subjects and tissue collection

Human reproductive tissues were obtained from individuals with genetically confirmed Helsmoortel-Van der Aa syndrome and non-syndromic unaffected control donors. Human tissue analyses were exploratory and aimed to provide translational validation of molecular findings identified in murine models. Given the rarity of Helsmoortel-Van der Aa syndrome and the limited availability of reproductive tissues, all available patient-derived specimens and matched control tissues were included. The study was approved by the Ethics Committee of the University Hospital Antwerp (UZA), Antwerp, Belgium, and registered under the Belgian Registration Number B300201627322. Control tissues were obtained through the Erasmus MC Tissue Bank (Rotterdam, the Netherlands) under approval code FO-WSB-023. Tissue collection, processing and used complied with institutional ethical guidelines and the Declaration of Helsinki.

#### Human testicular tissue

Control testicular tissue was obtained from three adolescent male donors aged 14-15 years through the Erasmus MC Tissue Bank, Department of Pathology, Erasmus University Medical Centre, Rotterdam, the Netherlands. Tissue originated from therapeutic resections performed for non-malignant clinical indications. Histopathological assessment confirmed the absence of neoplastic abnormalities. One specimen was obtained following unilateral left-sided testicular resection without evidence of malignancy, a second following right-sided testicular resection in the context of contralateral testicular atrophy, and a third following unilateral left-sided testicular resection without histopathological abnormalities. Only tissues classified as histologically normal by certified pathologists and devoid of malignant or overt inflammatory pathology were included as controls. Testicular tissue from one male individual with genetically confirmed Helsmoortel-Van der Aa syndrome was also analysed. The individual carried the *de novo* heterozygous *ADNP* frameshift variant c.1676dupA (p.His559Glnfs*3). Detailed clinical phenotyping has been reported previously (35). Tissue was obtained during autopsy following death at 6 years of age at the Clinical Department of Pathology and Cytology, University Hospital Centre Zagreb, Zagreb, Croatia. The post-mortem interval was approximately 35 h. Tissue processing and sectioning were subsequently performed by the Department of Neurology, Laboratory of Experimental Neurology, AHEPA University Hospital, Aristotle University of Thessaloniki, Thessaloniki, Greece. Tissue was divided for molecular and histological analyses, with molecular samples snap frozen in liquid nitrogen and stored at −80°C.

#### Human uterine tissue

Control uterine tissue was obtained from three adult female donors aged 30-38 years through the Erasmus MC Tissue Bank, Department of Pathology, Erasmus University Medical Centre, Rotterdam, the Netherlands. Tissues originated from surgical procedures performed for benign clinical indications and were confirmed to be free of malignancy or major structural abnormalities by histopathological assessment. One control sample was obtained from uterine resection tissue with cervical intraepithelial neoplasia grade 1 (CIN -1) and benign pelvic lymph node excision without evidence of malignancy. A second sample originated from pelvic soft tissue excision and uterine resection without pathological abnormalities. The third specimen consisted of uterus, tuba, and cervix tissue without histopathological abnormalities. Uterine tissue from one female individual with genetically confirmed Helsmoortel-Van der Aa syndrome was analysed. The individual carried the *de novo* heterozygous *ADNP* nonsense variant c.1211C>A (p.Ser404*). Her initial clinical characterisation was previously reported as patient II (6), with subsequent endocrine and reproductive features reported previously (10). Uterine tissue was obtained at 18 years of age and processed for molecular and histological analyses. Tissue processing was performed by the Department of Pathology, Erasmus University Medical Centre, Rotterdam, the Netherlands. Tissue for molecular analysis was snap frozen in liquid nitrogen and stored at −80°C, whereas parallel tissue fragments were fixed for histological analyses.

#### Human molecular analyses

Human testicular and uterine tissues were analysed by RT-qPCR, immunohistochemistry and immunoblotting to assess the translational relevance of selected murine findings. For RT-qPCR, total RNA was extracted using the RNeasy Mini Kit, and cDNA synthesis and quantitative PCR were performed as described above. Human gene expression was normalised to *TBP*, *RPL13A* and *RPLP0*. Representative genes identified in the murine transcriptomic analyses were assessed, including *ADH1C* and *TSPAN8* in testicular tissue and *CNN1* and *TGFBR3L* in uterine tissue. The ADNP-specific transcript assays were performed using previously published assays designed to distinguish total transcript abundance from transcripts retaining or spanning the deleted region (21). Statistical analyses were performed in GraphPad Prism (version 10.6.1) using a Welch’s t-test.

### Immunohistochemistry on human reproductive tissues

Formalin-fixed paraffin-embedded human testicular and uterine tissues were sectioned at 5 µm thickness using an Epredia HM 340E Electronic Rotary Microtome (Fisher Scientific; 10076358) and mounted on Superfrost Plus microscope slides (Avantor; 631-0108). Immunohistochemistry was performed as previously described (21). Briefly, sections were deparaffinised, rehydrated, and subjected to heat-mediated antigen retrieval using Tris/EDTA buffer (pH 9.0) prior to antibody incubation. Sections were incubated overnight at 4°C with primary antibodies against CNN1 (Abcam; RRID:AB_2291941) and TSPAN8 (Abcam; RRID: AB_3751514), diluted 1:250 in blocking solution. Following washing steps, sections were incubated with appropriate fluorophore-conjugated secondary antibodies. Nuclear counterstaining was performed using DAPI. Images were acquired using a Leica fluorescence microscopy under identical exposure conditions across samples. Protein intensity was quantified using ImageJ. Statistical analyses were performed in GraphPad Prism (version 10.6.1) using a Welch’s t-test.

### Clinical cohort re-analysis of Helsmoortel-Van der Aa syndrome

Clinical data from individuals with genetically confirmed Helsmoortel-Van der Aa syndrome were retrospectively re-analysed from a previously published international multicentre cohort comprising 144 individuals with pathogenic *ADNP* variants. Detailed cohort assembly, clinical phenotyping, variant classification and primary clinical characterisation have been reported previously (11). For the present analysis, the previously curated dataset was re-evaluated to investigate potential sex-dependent differences in clinical and developmental features. The analysed cohort comprised 129 individuals with available sex-stratified data, including 69 males and 60 females. Clinical features were coded as categorical variables according to their presence or absence.

Developmental measures, including age at first words and age at independent walking, were analysed as continuous variables. For categorical clinical features, associations between sex and the presence of individual features were assessed using Fisher’s exact test. Continuous developmental variables were compared between male and female individuals using Student’s t-tests. Given the broad range of clinical features assessed, false discovery rate correction was applied to the categorical and continuous clinical comparisons. q-values were calculated using the *qvalue* package in *R*. An FDR-adjusted q < 0.05 was considered statistically significant. Statistical analyses were performed in *R* (version 4.3.1), and data visualisation was performed using GraphPad Prism (version 10.6.1).

### Statistical analysis

Unless otherwise indicated, data are presented as mean ± standard deviation (s.d.). Statistical analyses were performed using GraphPad Prism (version 10.6.1) or *R* (version 4.3.1), as appropriate. For animal molecular experiments, statistical tests were selected according to the experimental design and are specified in the corresponding Methods section. For factorial experiments involving genotype and treatment, two-way ANOVA was used, followed by the specified multiple-comparisons procedure. Welch’s t-test was used for two-group comparisons when unequal variances were considered appropriate. For genome-wide DNA methylation analysis, statistical models and tests were selected according to the experimental design and are described in the corresponding DNA methylation Methods section. Differentially methylated features were identified using an FDR-adjusted p < 0.05 together with a biological effect-size threshold of |Δβ| ≥ 0.1. Multiple testing was controlled using the Benjamini-Hochberg false discovery rate (FDR) procedure. For transcriptome-wide RNA-sequencing analyses, differential expression was assessed using the DESeq2 Wald test for specified genotype comparisons and the DESeq2 likelihood ratio test (LRT) for genotype × treatment interaction effects. Genes were considered differentially expressed at an FDR-adjusted p < 0.05. For pathway and functional enrichment analyses, an adjusted p < 0.05 was considered statistically significant. For overlap analyses between male and female methylation or transcriptomic signatures, Fisher’s exact test was used after restricting the background universe to genes tested in both sexes. Jaccard indices and fold enrichment were calculated to quantify the magnitude of overlap. For the clinical cohort, Fisher’s exact test was used for categorical variables and unpaired, two-tailed Student’s t-test for continuous variables, with FDR correction applied to the clinical feature comparisons. For analyses not otherwise specified, two-sided p < 0.05 was considered statistically significant. P-values and FDR-adjusted values are reported as indicated in the corresponding Results and figure legends.

### Experimental Flow-Chart

A graphical illustration of the experimental design and experimental procedures conducted within this study (figure 1).

**Figure 1.**
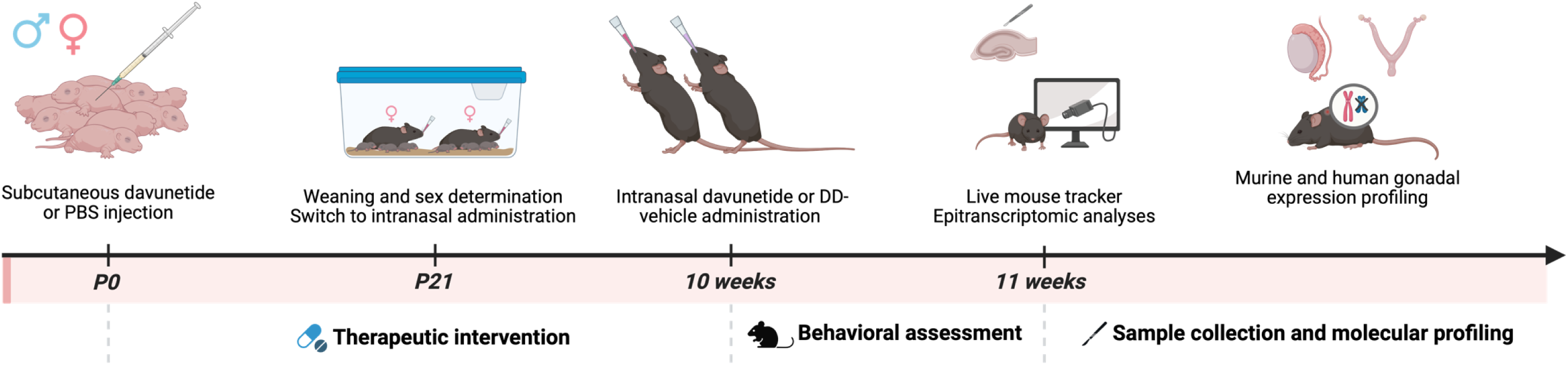
Experimental workflow for behavioural, molecular, and translational analyses in male and female *Adnp* HET mice. Schematic overview of the experimental design used to investigate sex-dependent behavioural and molecular effects in *Adnp* HET mice and WT littermate controls. Litters were randomly assigned at birth (postnatal day 0, P0) to receive either NAP (davunetide) or DD-vehicle prior to genotyping and sex determination at weaning (P21). During early postnatal development (P0-P21), mice received daily subcutaneous injections of NAP or DD-vehicle, followed by daily intranasal administration after weaning until sacrifice. Behavioural phenotyping was conducted at 10 weeks of age using the Live Mouse Tracker (LMT) platform, enabling continuous 24-hour automated behavioural assessment under DD-vehicle conditions and evaluation of behavioural responsiveness to NAP (davunetide). At 11 weeks, hippocampal and gonadal tissues (testis and uterus) were collected for downstream molecular analyses, including immunoblotting, RT-qPCR, DNA methylation profiling, and bulk RNA sequencing. These data were integrated with publicly available RNA sequencing datasets from the four-core genotype (4CG) mouse model to assess sex-dependent and hormone-dependent *Adnp* gene expression. Finally, murine findings were translated to human gonadal tissue analyses and clinical cohort data from individuals with Helsmoortel-Van der Aa syndrome.

## Results

### Global behavioural organisation is altered by Adnp deficiency

To determine whether *Adnp* deficiency alters spontaneous behavioural organisation in a sex-dependent manner, male and female WT and HET mice were continuously monitored for 24 hours using the LMT platform (22). Behavioural recordings were performed in mixed-genotype social groups within the same cage, generating 33 individual and social behavioural parameters for each animal.

We first performed an unsupervised principal component analysis (PCA) including all male and female WT and HET mice maintained under DD-vehicle conditions. The first two principal components explained 39.2% of the total behavioural variance and showed genotype-associated separation in behavioural organisation across both sexes (figure 2A). Although WT males and females occupied distinct behavioural spaces, *Adnp* deficiency displaced behavioural profiles in both sexes, with female HET mice showing the greatest visual separation from their WT counterparts. To quantify genotype- or sex-associated differences in global behavioural organisation, individual principal component 1 (PC1) scores were compared between groups. Both male (male WT: 0.22 ± 0.26; male HET: -0.25 ± 0.35; 95% CI: 0.27 to 0.66; two-way ANOVA, p < 0.0001) and female (female WT: 0.26 ± 0.34; female HET: -0.16 ± 0.22; 95% CI: 0.22 to 0.62; two-way ANOVA, p < 0.0001) HET mice exhibited significantly different PC1 values compared with their respective WT littermates, whereas no significant global differences were observed between WT males and females (male WT: 0.22 ± 0.26; female WT: 0.26 ± 0.34; 95% CI: -0.24 to 0.15; two-way ANOVA, p > 0.9999) or between male and female HET mice (male HET: -0.25 ± 0.35; female HET: -0.16 ± 0.22; 95% CI: -0.29 to 0.10; two-way ANOVA, p > 0.9999) (figure 2B).

**Figure 2.**
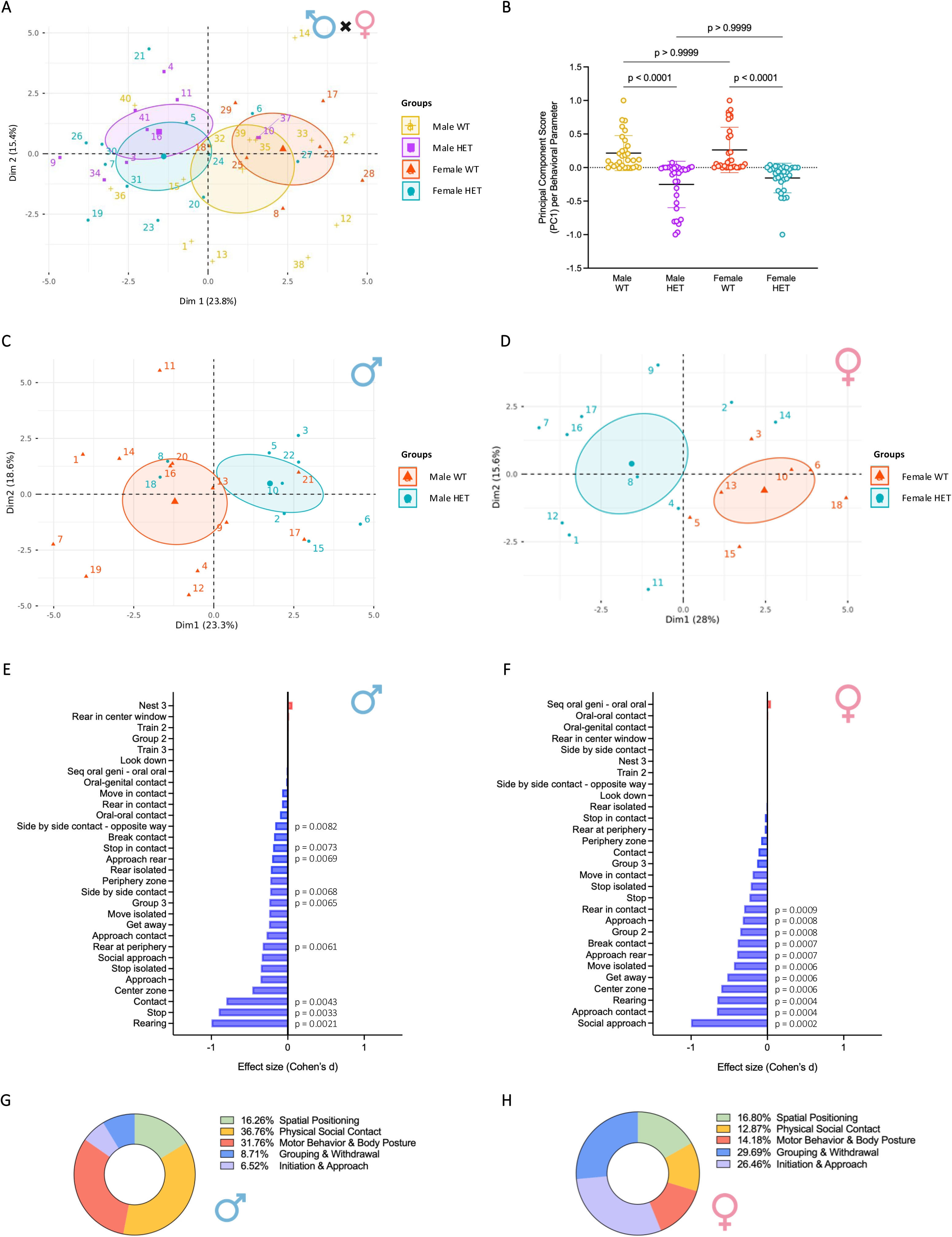
Adnp deficiency is associated with sex-specific behavioural organisation in male and female mice following automated 24-hour behavioural phenotyping. **(A)** Principal component analysis (PCA) of all 33 behavioural parameters obtained using the LMT platform from male and female WT and HET mice receiving DD-vehicle solution. The first two principal components explain 39.2% of the total behavioural variance. Each point represents one individual mouse, and ellipses indicate the 95% confidence interval for each group. **(B)** Comparison of principal component 1 (PC1) scores demonstrating significant genotype-associated differences between HET and WT mice in both males and females, with no significant differences between sexes. Each dot represents one behavioural parameter (n = 33 behavioural parameters per group). Statistical analysis was performed using two-way ANOVA with genotype and sex as factors, followed by Tukey’s multiple comparisons test. Data are presented as mean ± s.d. **(C,D)** Sex-stratified PCA showing genotype-associated behavioural organisation between WT and HET mice in males **(C)** and females **(D)**. **(E,F)** Cohen’s *d* effect sizes for the 33 behavioural variables in males **(E)** and females **(F)**. Positive effect sizes indicate behaviours increased in HET mice relative to WT mice, whereas negative effect sizes indicate behaviours decreased in HET mice relative to WT mice. Behavioural variables were analysed using linear mixed-effects models (LMM) with genotype as a fixed effect and cage identity as a random intercept. **(G,H)** Relative contribution of the five predefined LMT behavioural domains to the overall genotype-associated behavioural phenotype in males **(G)** and females **(H)**. Domain contributions were calculated by summing the absolute genotype-associated effect sizes (Cohen’s *d*) of all behavioural variables assigned to each domain and expressing these values as a percentage of the total summed absolute effect size across all behavioural variables. The five behavioural domains comprised physical social contact, motor behaviour and body posture, spatial positioning, grouping and withdrawal, and initiation and approach.

### Finer-scale behavioural features differ between male and female Adnp heterozygous mice

To identify behavioural features underlying these global similarities, males and females were subsequently analysed separately. PCA demonstrated clear genotype-dependent separation between WT and HET mice within each sex (males: LMM, padj. = 0.0377 and females: LMM, padj. = 0.0189) (figure 2C,D), indicating that although the magnitude of behavioural disruption was comparable, the behavioural architecture differed between males and females. Effect size analysis of the 33 behavioural variables demonstrated distinct sex-specific behavioural signatures (figure 2E,F). In males, the largest genotype-associated effect sizes were observed for reduced rearing (Cohen’s *d*: -1; LMM, p = 0.0021), stop contact (Cohen’s *d*: -0.90; LMM, p = 0.0033), contact (Cohen’s *d*: -0.80; LMM, p = 0.0043), rear at periphery (Cohen’s *d*: -0.33; LMM, p = 0.0061), group of three formation (Cohen’s *d*: -0.24; LMM, p = 0.0065), side-by-side contact (Cohen’s *d*: -0.23; LMM, p = 0.0068), approach rear (Cohen’s *d*: -0.20; LMM, p = 0.0069), stop in contact (Cohen’s *d*: -0.19; LMM, p = 0.0073), and side-by-side contact in opposite orientation (Cohen’s *d*: -0.16; LMM, p = 0.0082). In females, the strongest genotype-associated effects involved social approach (Cohen’s *d*: -1; LMM, p = 0.0002), approach contact (Cohen’s *d*: -0.66; LMM, p = 0.0004), rearing (Cohen’s *d*: -0.65; LMM, p = 0.0004), centre-zone occupancy (Cohen’s *d*: -0.60; LMM, p = 0.0006), get away (Cohen’s *d*: -0.53; LMM, p = 0.0006), move isolated (Cohen’s *d*: -0.44; LMM, p = 0.0006), approach rear (Cohen’s *d*: -0.40; LMM, p = 0.0007), break contact (Cohen’s *d*: -0.39; LMM, p = 0.0007), group of two formation (Cohen’s *d*: -0.36; LMM, p = 0.0008), approach (Cohen’s *d*: -0.32; LMM, p = 0.0008), and rear in contact (Cohen’s *d*: -0.30; LMM, p = 0.0009). Collectively, these behavioural variables indicate that male HET mice are primarily characterised by alterations in exploratory activity, physical social contact, and group-level interactions, whereas female HET mice exhibit greater disruption of social investigation, social approach, locomotor activity, centre -zone exploration, and smaller group dynamics.

To facilitate biological interpretation, these behavioural variables were subsequently grouped into predefined behavioural domains of the LMT framework. For each domain, the absolute genotype-associated effect sizes of all constituent behaviours were summed and expressed as a percentage of the total summed effect size across all behavioural variables. In males, the genotype-dependent behavioural differences were predominantly explained by physical social contact (36.76%), motor behaviour and body posture (31.76%), and spatial positioning (16.26%), whereas grouping and withdrawal (8.71%) and initiation and approach (6.52%) contributed less prominently (figure 2G). Conversely, female behavioural alterations were dominated by grouping and withdrawal (29.69%) and initiation and approach (26.46%), with comparatively smaller contributions from spatial positioning (16.80%), motor behaviour and body posture (14.18%), and physical social contact (12.87%) (figure 2H).

Collectively, these findings indicate that *Adnp* deficiency produces a comparable degree of global behavioural alteration in male and female mice, while revealing distinct finer-scale, sex-specific behavioural profiles underlying this phenotype. Male behavioural alterations were primarily characterised by exploratory and contact-associated behaviours, whereas females exhibited proportionally greater disruption of social organisation and interaction.

### Chronic NAP administration partially normalises behavioural abnormalities in both sexes

Because the ADNP-derived octapeptide NAP (davunetide) has been investigated as a potential therapeutic candidate for Helsmoortel-Van der Aa syndrome (23), we next assessed whether long-term treatment from birth to 10 weeks of age could ameliorate behavioural abnormalities identified using the LMT platform.

In male mice, PCA demonstrated that DD-vehicle HET animals formed the most distinct behavioural cluster, whereas NAP-treated HET mice shifted towards the WT behavioural space (LMM, padj. = 0.0225), indicating partial normalisation of behavioural organisation (figure 3A). A comparable pattern was observed in females (LMM, padj. = 0.0283), although the behavioural shift towards WT profiles appeared less pronounced, with NAP-treated HET mice remaining more separated from WT animals compared with males (figure 3B). Effect size analysis further identified the behavioural variables associated with NAP-mediated behavioural improvement (figure 3C,D). In male HET mice, NAP treatment predominantly increased stop in contact (Cohen’s *d*: 1; LMM, p = 0.011), contact (Cohen’s *d*: 0.94; LMM, p = 0.014), social approach (Cohen’s *d*: 0.88; LMM, p = 0.018), nose-anogenital contact (Cohen’s *d*: 0.52; LMM, p = 0.024), sequential oral-genital/oral interactions (Cohen’s *d*: 0.52; LMM, p = 0.028), and group of three formation (Cohen’s *d*: 0.49; LMM, p = 0.033). In female HET mice, treatment primarily affected approach contact (Cohen’s *d*: 1; LMM, p = 0.013), centre-zone occupancy (Cohen’s *d*: 0.93; LMM, p = 0.019), break contact (Cohen’s *d*: 0.84; LMM, p = 0.026), approach (Cohen’s *d*: 0.83; LMM, p = 0.033), and get away behaviours (Cohen’s *d*: 0.66; LMM, p = 0.043). These findings indicate that NAP predominantly improves social interaction patterns in both sexes, although the specific behavioural features associated with improvement differ between males and females.

**Figure 3.**
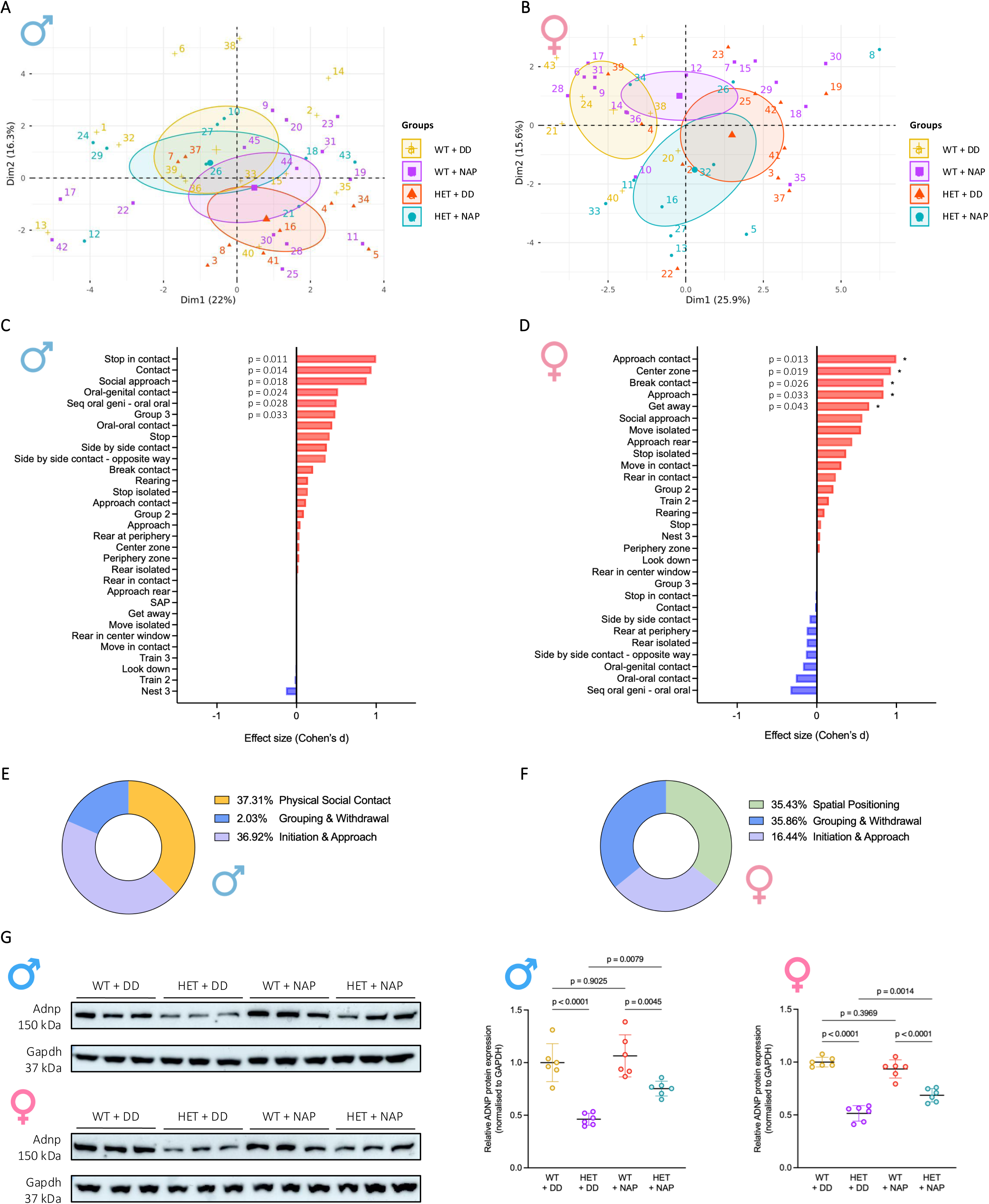
Chronic NAP treatment partially normalises behavioural organisation and increases hippocampal ADNP protein abundance in HET mice. (A,B) Principal component analyses of behavioural profiles in male **(A)** and female **(B)** WT and HET mice following chronic NAP or DD-vehicle treatment. Ellipses represent the 95% confidence intervals for each group. Statistical comparison of behavioural profiles was performed using LMMs based on individual PC scores. **(C,D)** Cohen’s *d* effect sizes illustrating behavioural variables associated with NAP treatment in males **(C)** and females **(D)**. Positive effect sizes indicate behaviours increased following NAP treatment, whereas negative effect sizes indicate behaviours decreased relative to DD-vehicle-treated HET mice. Behavioural variables were analysed using LMMs with treatment as a fixed effect and cage identity as a random intercept. **(E,F)** Relative contribution of the five predefined LMT behavioural domains associated with NAP treatment in males **(E)** and females **(F)**. Domain contributions were calculated by summing the absolute treatment-associated effect sizes (Cohen’s *d*) of all behavioural variables assigned to each domain and expressing these values as a percentage of the total summed absolute effect size across all behavioural variables. **(G)** Representative immunoblots showing hippocampal ADNP protein abundance in WT and HET mice following DD-vehicle or NAP treatment, with GAPDH used as a loading control. Quantification demonstrates reduced hippocampal ADNP protein abundance in HET mice under DD-vehicle conditions and increased ADNP protein abundance following NAP treatment in HET mice of both sexes. Statistical analyses were performed using two-way ANOVA followed by Tukey’s multiple comparisons test. Data are presented as mean ± s.d. (n = 6 per genotype and treatment group).

Behavioural effect size analysis further demonstrated that NAP treatment predominantly affected social behavioural features in both sexes, although the behavioural domains showing the largest relative changes differed. In males, NAP-associated behavioural changes were predominantly observed in physical social contact (37.31%), followed by initiation and approach (36.92%), and grouping and withdrawal (2.03%) (figure 3E). In females, NAP-associated changes were primarily observed in grouping and withdrawal (35.86%), spatial positioning (35.43%), and initiation and approach (16.44%) (figure 3F). Together, these findings indicate that NAP partially normalises social behavioural organisation in both sexes, while the specific behavioural domains affected differ between males and females.

To determine whether behavioural improvement was accompanied by changes in overall somatic growth, body weight was assessed throughout the treatment period (additional file S4). Under DD-vehicle conditions, both male (male WT + DD: 28.04 g ± 1.57; male HET + DD: 23.59 ± 2.34; 95% CI: 2.81 to 6.10; two-way ANOVA, p < 0.0001) and female (female WT + DD: 20.45 g ± 1.46; female HET + DD: 18.32 ± 1.69; 95% CI: 1.16 to 3.31; two-way ANOVA, p < 0.0001) HET mice exhibited significantly reduced body weight compared with WT littermates, consistent with findings in individuals with Helsmoortel-Van der Aa syndrome (10). Chronic NAP administration did not significantly alter body weight in either genotype (male WT + DD: 28.04 g ± 1.57; male WT + NAP: 27.93 ± 1.92; 95% CI: -1.37 to 1.81; two-way ANOVA, p = 0.9940; male HET + DD: 23.59 g ± 2.34; male HET + NAP: 24.07 ± 1.88; 95% CI: -2.64 to 0.88; two-way ANOVA, p = 0.6009; female WT + DD: 20.45 g ± 1.46; female WT + NAP: 20.69 ± 1.06; 95% CI: -1.69 to 0.45; two-way ANOVA, p = 0.4648; female HET + DD: 18.32 g ± 1.69; female HET + NAP: 18.95 ± 1.10; 95% CI: -2.00 to 0.14; two-way ANOVA, p = 0.1131), indicating that behavioural changes occurred independently of detectable changes in overall body size or growth trajectory.

We subsequently evaluated whether behavioural improvement was associated with changes in ADNP protein abundance, a molecular target of NAP-mediated activity (39). ADNP protein expression was quantified in hippocampal lysates by immunoblotting (figure 3G). As expected, hippocampal ADNP protein abundance was reduced by approximately 50% in DD-vehicle-treated HET mice of both sexes compared with WT littermates (male WT + DD: 1.00 ± 0.17; male HET + DD: 0.46 ± 0.06; 95% CI: -0.31 to 0.76; two-way ANOVA, p < 0.0001; female WT + DD: 1.00 ± 0.05; female HET + DD: 0.51 ± 0.07; 95% CI: 0.38 to 0.59; two-way ANOVA, p < 0.0001). NAP treatment did not alter ADNP protein expression in WT mice (male WT + DD: 1.00 ± 0.17; male WT + NAP: 1.06 ± 0.20; 95% CI: -0.29 to 0.16; two-way ANOVA, p = 0.9025; female WT + DD: 1.00 ± 0.05; female WT + NAP: 0.94 ± 0.09; 95% CI: -0.04 to 0.17; two-way ANOVA, p = 0.3969). In contrast, NAP significantly increased hippocampal ADNP protein abundance in HET animals of both sexes relative to genotype-matched DD-vehicle controls (male HET + DD: 0.46 ± 0.06; male HET + NAP: 0.75 ± 0.07; 95% CI: -0.52 to -0.07; two-way ANOVA, p = 0.0079; female HET + DD: 0.51 ± 0.07; female HET + NAP: 0.69 ± 0.06; 95% CI: -0.28 to -0.06; two-way ANOVA, p = 0.0014). Although the relative increase appeared more pronounced in males, NAP treatment increased ADNP protein abundance in both male and female HET mice, consistent with a shared molecular response to treatment across sexes.

To determine whether NAP-mediated increases in ADNP protein abundance were due to altered expression of the mutant *Adnp* allele, allele-specific RT-qPCR was performed using primers spanning the CRISPR-generated frameshift variant (additional file S5). Mutant transcripts were detected exclusively in HET mice (male WT + DD: 0.00 ± 0.00; male HET + DD: 1.01 ± 0.16; 95% CI: -1.18 to -0.84; two-way ANOVA, p < 0.0001; female WT + DD: 0.01 ± 0.01; female HET + DD: 1.02 ± 0.23; 95% CI: -1.23 to -0.80; two-way ANOVA, p < 0.0001) and remained unchanged following NAP treatment in either sex (male HET + DD: 1.01 ± 0.16; male HET + NAP: 0.95 ± 0.14; 95% CI: -0.11 to 0.23; two-way ANOVA, p = 0.7561; female HET + DD: 1.02 ± 0.23; female HET + NAP: 1.10 ± 0.28; 95% CI: -0.29 to 0.14; two-way ANOVA, p = 0.7818), demonstrating that NAP-associated increases in ADNP protein abundance were not accompanied by increased mutant *Adnp* transcript expression. Consistent with protein analyses, global *Adnp* transcript levels were reduced in HET mice relative to WT controls (male WT + DD: 1.01 ± 0.15; male HET + DD: 0.72 ± 0.11; 95% CI: 0.12 to 0.46; two-way ANOVA, p = 0.0002; female WT + DD: 1.02 ± 0.20; female HET + DD: 0.63 ± 0.10; 95% CI: 0.17 to 0.60; two-way ANOVA, p = 0.0001). NAP treatment produced a non-significant trend towards increased total *Adnp* transcript abundance in female HET mice (female HET + DD: 0.63 ± 0.10; female HET + NAP: 0.84 ± 0.07; 95% CI: -0.43 to 0.01; two-way ANOVA, p = 0.0623), whereas no transcriptional response was observed in male HET mice (male HET + DD: 0.72 ± 0.01; male HET + NAP: 0.73 ± 0.12; 95% CI: -0.18 to 0.16; two-way ANOVA, p = 0.9970).

### Sex-dependent DNA methylation and gene expression in Adnp-deficient mice

To determine whether the finer-scale sex-dependent behavioural phenotypes observed in *Adnp*-deficient mice were accompanied by corresponding molecular alterations, we performed genome-wide DNA methylation profiling of hippocampal tissue from WT and HET male and female mice. Linear mixed models incorporating sex or genotype as explanatory variables were applied to identify sex- and genotype-associated methylation changes, respectively. These analyses identified 334 sex-associated CpG probes (LMM, FDR-adjusted p < 0.0001) (additional file S6) and 3,483 genotype-associated CpG probes (LMM, FDR-adjusted p < 0.01) (additional file S7), demonstrating that the *Adnp* genotype represents the predominant determinant of hippocampal methylation variation, while sex contributes an additional layer of molecular regulation.

Given the established contribution of sex to neurobiological regulation (40), we first characterised sex-associated methylation differences. Differentially methylated probes were defined according to the male versus female comparison, such that positive Δβ values indicate higher methylation in males and negative Δβ values indicate lower methylation in males relative to females. Comparison of male and female mice within each genotype identified 918 unique sex-associated CpGs (78.4%), whereas the corresponding WT comparison identified 178 unique CpGs (15.2%). A further 75 CpGs (6.4%) were shared between both genotype comparisons (figure 4A). Thus, although sex-associated methylation differences were detectable in both genotypes, the substantial larger number of sex-associated CpGs in HET mice suggests that loss of *Adnp* may modify sex-dependent regulation of the hippocampal methylome.

**Figure 4.**
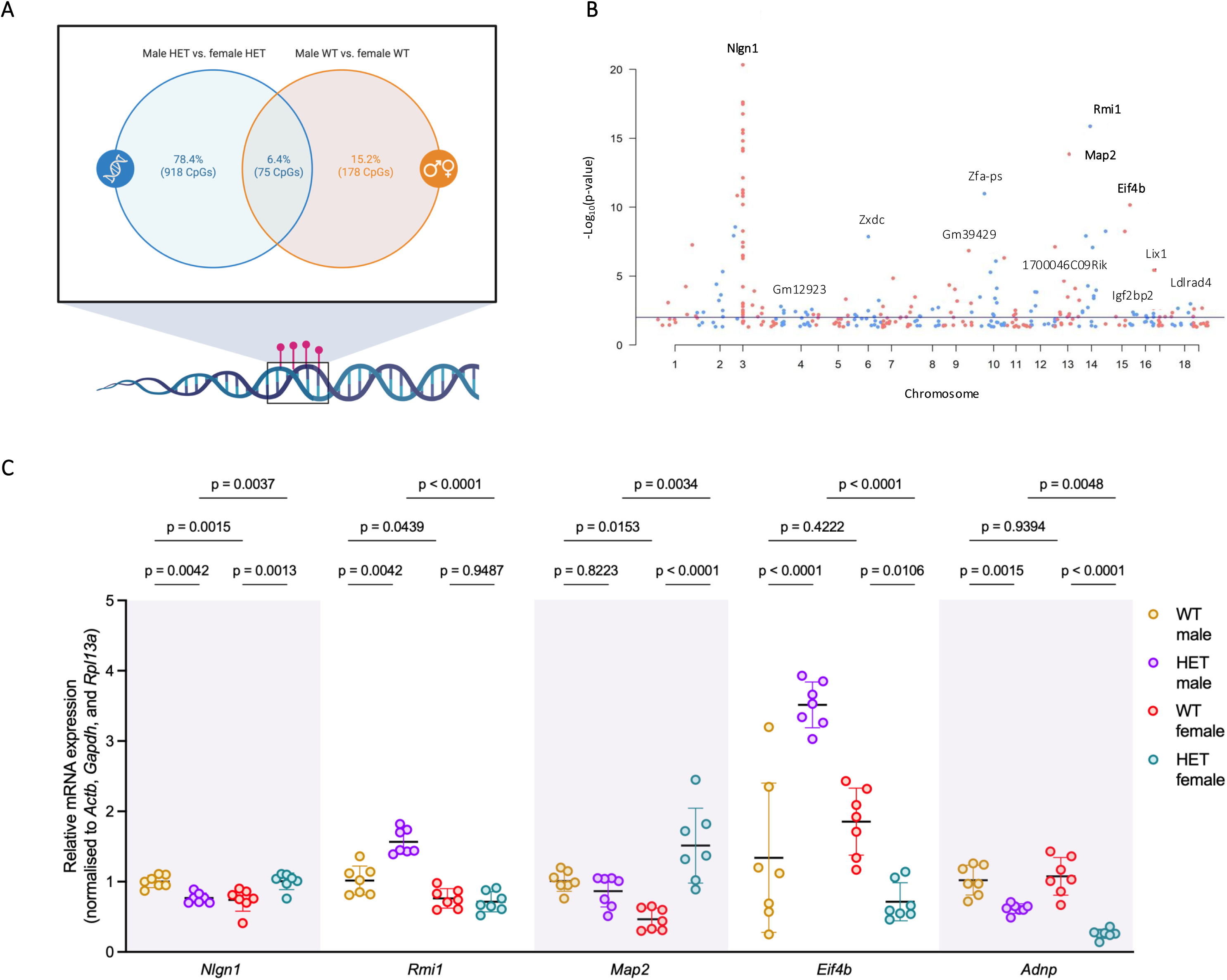
Sex-associated DNA methylation and transcriptional alterations in the hippocampus of *Adnp*-deficient mice. **(A)** Venn diagram showing overlap between sex-associated differentially methylated CpG probes identified by comparing male and female mice within each genotype. Differential methylation was calculated as male minus female, such that positive Δβ values indicate higher methylation in males and negative Δβ values indicate lower methylation in males relative to females. The majority of sex-associated CpGs were detected in HET mice, indicating enhanced sexually dimorphic methylation following *Adnp* deficiency. **(B)** Manhattan plot displaying sex-associated CpG probes identified by linear mixed modelling with sex as an explanatory variable. The x-axis indicates chromosomal position and the y-axis indicates −log10(FDR-adjusted p-value). Major sex-associated methylation loci are indicated. **(C)** Relative mRNA expression of selected methylation-associated genes (*Nlgn1*, *Rmi1*, *Map2*, *Eif4b*, and *Adnp*) measured by RT-qPCR in hippocampal tissue from WT and HET male and female mice. Statistical analysis was performed using two-way ANOVA with genotype and sex as factors followed by Tukey’s multiple comparisons test. Data are presented as mean ± s.d. (n = 7 per genotype and treatment group).

To further define sex-associated methylation changes, the sex-based linear mixed model identified 334 significant CpG probes (LMM, FDR-adjusted p < 0.0001), of which 212 probes corresponding to 190 unique CpG positions mapped to 171 annotated genes (figure 4B). The strongest sex-associated DNA methylation differences were observed at *Nlgn1*, where 38 significant probes (16 unique CpG positions; chr3:25,520,123–25,674,039) were identified, including 32 probes reaching a more stringent significance threshold (LMM, FDR-adjusted p < 0.00001). Among probes exceeding the predefined biological effect size threshold (|Δβ| > 0.1), *Nlgn1* exhibited higher methylation in males compared with females (mean Δβ = +0.11), with the strongest difference observed within a concordant six-probe cluster (chr3:25,531,667; mean Δβ = +0.39). Additional sex-associated methylation changes were identified at *Rmi1* (Δβ = −0.14), *Map2* (Δβ = −0.12), and *Eif4b* (Δβ = −0.15) among other loci (additional file S8).

To determine whether sex-associated methylation differences were reflected at the transcriptional level, five candidate genes (*Nlgn1*, *Rmi1*, *Map2*, *Eif4b*, and *Adnp*) were selected for RT-qPCR validation based on methylation significance or biological relevance to ADNP-associated pathways (13,18,41,42). RT-qPCR analysis demonstrated that several methylation-associated loci displayed sex-dependent transcriptional responses following *Adnp* deficiency (figure 4C). *Nlgn1* expression showed a sex-dependent response to *Andp* deficiency, with reduced expression in HET males compared with HET females (male HET: 0.77 ± 0.07; female HET: 1.01 ± 0.12; 95% CI: -0.41 to -0.07; two-way ANOVA, p = 0.0037), consistent with the hypermethylation identified in males. *Rmi1* expression was increased in HET males compared with HET females (male HET: 1.57 ± 0.18; female HET: 0.71 ± 0.15; 95% CI: 0.60 to 1.10; two-way ANOVA, p < 0.0001), in line with the hypomethylation observed in males at this locus. *Map2* expression demonstrated a female-specific response to *Adnp* deficiency (female WT: 0.47 ± 0.15; female HET: 1.51 ± 0.54; 95% CI: -1.50 to -0.59; two-way ANOVA, p < 0.0001), together with increased expression in female HET compared with male HET mice (male HET: 0.87 ± 0.23; female HET: 1.51 ± 0.54; 95% CI: -1.10 to -1.19; two-way ANOVA, p = 0.0034), contrasting the predicted hypomethylation of *Map2* in males. Moreover, *Eif4b* and *Adnp* displayed genotype-associated transcriptional alterations across sexes. Eif4b expression was increased in male HET mice compared with male WT controls (male WT: 1.34 ± 1.08; male HET: 3.51 ± 0.34; 95% CI: -3.09 to -1.26; two-way ANOVA, p < 0.0001) and altered in female mice (female WT: 1.85 ± 0.45; HET: 0.71 ± 0.27; 95% CI: 0.23 to 2.05; two-way ANOVA, p = 0.0106) with additional sex-dependent differences between HET males and females (male HET: 3.51 ± 0.33; female HET: 0.71 ± 0.27; 95% CI: 1.89 to 3.71; two-way ANOVA, p < 0.0001). *Adnp* expression was reduced in HET males (male WT: 1.02 ± 0.21; male HET: 0.62 ± 0.07; 95% CI: 0.14 to 0.66; two-way ANOVA, p = 0.0015) and HET females (female WT: 1.08 ± 0.27; female HET: 0.26 ± 0.06; 95% CI: 0.56 to 1.08; two-way ANOVA, p < 0.0001), with lower expression in female HET mice compared with male HET mice (male HET: 0.62 ± 0.07; female HET: 0.26 ± 0.06; 95% CI: 0.10 to 0.62; two-way ANOVA, p = 0.0048).

Collectively, these findings demonstrate that *Adnp* deficiency produces predominantly genotype-driven hippocampal methylation differences, accompanied by a significant sex-associated molecular component. Sex-dependent methylation signatures are reflected by transcriptional alterations in biologically relevant ADNP-associated genes, providing a molecular framework for the sex-dependent behavioural phenotypes observed in *Adnp*-deficient mice.

### Genotype-dependent DNA methylation changes in Adnp-deficient mice

Following the identification of a predominant genotype-associated component of the hippocampal methylome, we next characterised the genome-wide DNA methylation changes associated with the *Adnp* frameshift variant within each sex. Unsupervised UMAP analysis of hippocampal methylation profiles showed genotype-associated separation in both males and females, with greater overlap between WT and HET profiles in females (figure 5A,B). Hierarchical clustering of differentially methylated CpG probes further separated male WT and HET mice according to genotype, whereas female mice showed a similar overall pattern, with two animals displaying less pronounced genotype-associated clustering (figure 5C,D). In males, 2,500 CpG probes met the predefined biological effect size threshold of |Δβ| > 0.1, comprising 184 hypermethylated (Δβ > 0.1) and 2,316 hypomethylated (Δβ < −0.1) probes (additional file S9). These probes were associated with 104 genes in the hypermethylated set and 988 protein-coding genes in the hypomethylated set, demonstrating a marked genome-wide shift towards hypomethylation in male HET mice. In females, 1,375 CpG probes were differentially methylated, including 16 hypermethylated and 1,359 CpG hypomethylated probes, associated with 9 and 652 protein-coding genes, respectively (additional file S10). Thus, female HET mice exhibited approximately 1.8-fold fewer differentially methylated probes than males, while both sexes showed striking predominance of hypomethylation, identifying widespread CpG hypomethylation as the principal methylation signature of *Adnp* deficiency.

**Figure 5.**
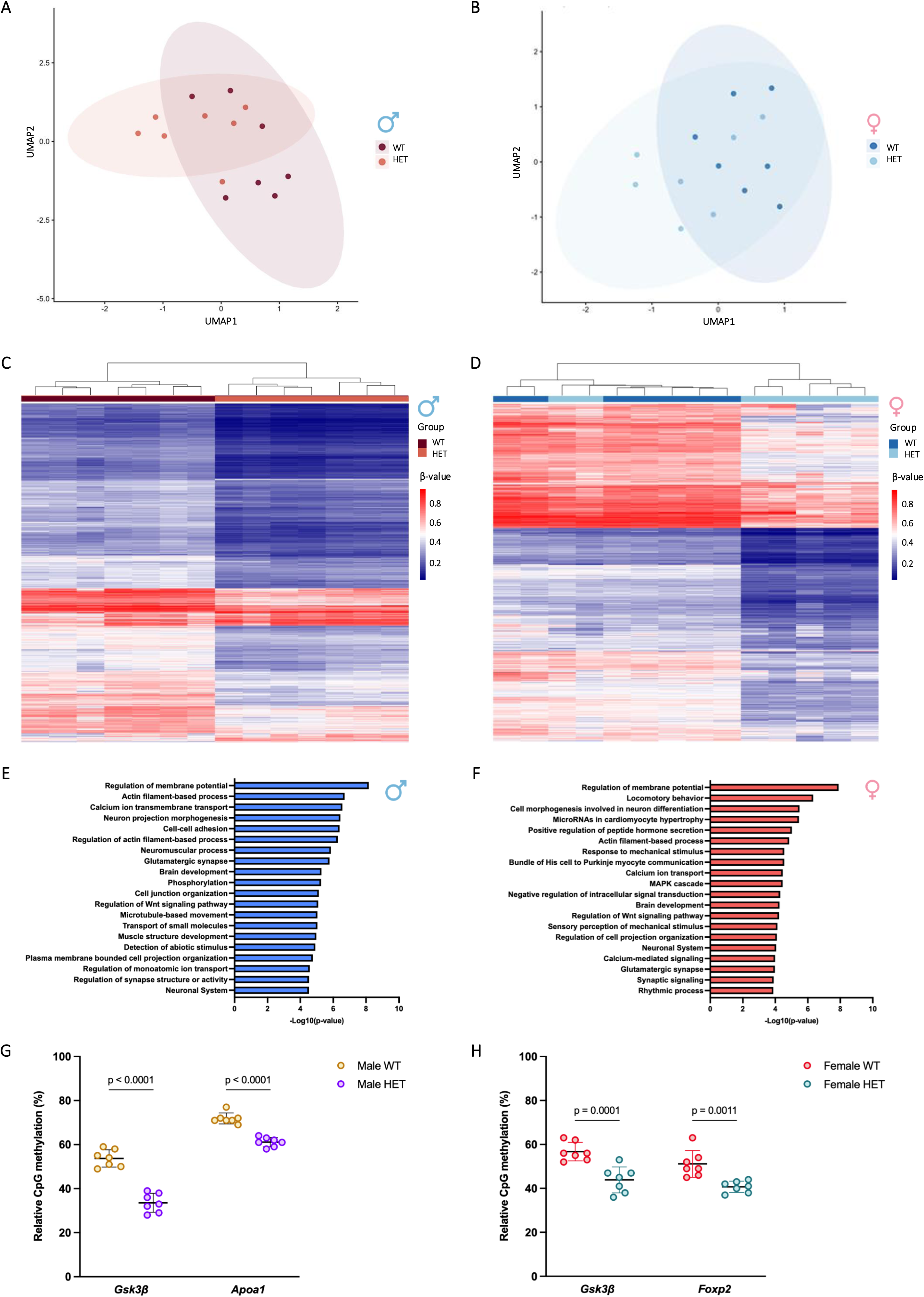
*Adnp* deficiency is associated with genome-wide hippocampal DNA hypomethylation in male and female mice. **(A,B)** UMAP visualisation of hippocampal DNA methylation profiles in male **(A)** and female **(B)** mice, showing genotype-associated separation of WT and HET methylation profiles. **(C,D)** Heatmaps showing genotype-associated DNA methylation profiles in male (C) and female (D) mice. CpG probes meeting the predefined biological effect size threshold of |Δβ| > 0.1 are shown as β-values across WT and HET animals. Hierarchical clustering separated male WT and HET samples by genotype, whereas female samples showed a similar overall pattern with two less pronounced genotype-associated profiles. Red and blue indicate relatively higher and lower DNA methylation, respectively. **(E,F)** Functional enrichment analysis of genes associated with genotype-dependent methylation changes in male **(E)** and female **(F)** mice, respectively, performed using Metascape. Shared enriched biological processes included regulation of membrane potential, actin filament-based processes, glutamatergic synapse, brain development and nervous system. **(G,H)** Targeted pyrosequencing validation of representative genotype-associated methylation changes in male **(G)** and female **(H)** mice. *Gsk3β* showed hypomethylation in both male and female HET mice compared with sex-matched WT littermates, whereas *Apoa1* and *Foxp2* showed hypomethylation identified in the male and female genotype-associated methylation signatures, respectively. Statistical analysis was performed using two-way ANOVA followed by Tukey’s multiple comparisons test. Data are presented as mean ± s.d. (n = 7 per genotype).

To quantify the convergence of genotype-associated methylation changes between sexes, we compared the genes associated with differentially methylated probes. Of 1,092 genes identified in males and 661 in females, 444 were shared between sexes, corresponding to 40.8% and 67.2% of the respective gene sets (Jaccard index = 34.0%). This overlap was substantially greater than expected by chance (13.1-fold enrichment; odds ratio = 63.10; 95% CI: 52.71 to 75.61; Fisher’s exact test, p < 10^-439^), demonstrating convergence of the genotype-associated methylation landscape despite the greater number of differentially methylated loci in males.

Functional enrichment analysis further supported this convergence (figure 5E,F). Five of the 20 most significantly enriched biological processes were shared between sexes, including regulation of membrane potential, actin filament-based processes, glutamatergic synaptic function, brain development, and neuronal system. These findings suggest that genotype-associated methylation changes in both sexes converge on biological processes relevant to neuronal function, synaptic signalling and cellular organisation.

We next validated representative genotype-associated methylation changes by targeted pyrosequencing (figure 5G,H), selecting *Gsk3β* as a shared ADNP-relevant target and *Apoa1* and *Foxp2* as representative male- and female-associated loci, respectively (20,43,44). Pyrosequencing confirmed the array-predicted hypomethylation of *Gsk3β* in male HET mice (male WT: 53.71 ± 3.64; male HET: 33.57 ± 4.32; 95% CI: 15.91 to 24.38; two-way ANOVA, p < 0.0001) and female HET mice (female WT: 56.71 ± 4.31; male HET: 43.86 ± 6.39; 95% CI: 6.60 to 19.12; two-way ANOVA, p = 0.0001) compared with sex-matched WT littermates. *Apoa1* was hypomethylated in male HET mice (male WT: 71.86 ± 2.73; male HET: 61.14 ± 2.27; 95% CI: 6.48 to 14.95; two-way ANOVA, p < 0.0001), whereas *Foxp2* was hypomethylation in female HET mice (female WT: 51.14 ± 5.70; female HET: 40.71 ± 2.06; 95% CI: 4.17 to 16.69; two-way ANOVA, p = 0.0011), confirming the direction of the genotype-associated methylation changes identified by the array. Collectively, these findings establish widespread, predominantly hypomethylating changes as the molecular signature of *Adnp* deficiency, with substantial convergence between sexes at the level of both affected genes and biological pathways.

The convergence of genotype-associated methylation changes on neuronal and actin filament-based processes in both sexes prompted us to examine whether postnatal treatment with the investigational ADNP-derived peptide NAP could modify the hippocampal methylation landscape (figure 1). We therefore performed genome-wide DNA methylation profiling of hippocampal tissue from male and female WT and HET mice treated with NAP or the DD-vehicle from postnatal treatment initiation until 10 weeks of age, using sex-matched vehicle-treated animals as controls.

To determine whether NAP treatment differentially affected the *Adnp* genotypes, we first fitted a linear mixed model incorporating genotype, treatment, and their interaction. No CpG probes showed a significant genotype × treatment interaction (LMM, FDR-adjusted p > 0.01) in either male or female mice, indicating that NAP treatment did not produce a genotype-dependent alteration of the hippocampal methylome (additional file S11). We therefore fitted a second model to evaluate treatment-associated methylation changes independent of the genotype, resulting in no genome-wide treatment-associated CpG methylation changes (LMM, FDR-adjusted p > 0.01) in either male or female mice (additional files S12). Consistent with these findings, unsupervised UMAP analysis and hierarchical clustering showed that methylation profiles remained primarily structured by genotype rather than treatment in both sexes (additional file S13, A-D).

Together, these analyses indicate that postnatal NAP treatment neither modifies the genotype-associated hippocampal methylation landscape nor induces a detectable genome-wide methylation response in WT or HET mice. Thus, the widespread hypomethylation associated with the *Adnp* frameshift variant is not measurably reversed by NAP treatment at the level of the hippocampal methylome.

### Sex-specific hippocampal transcriptomic alterations in Adnp-deficient mice and genotype-dependent responses to NAP treatment

Following the identification of predominantly genotype-associated DNA methylation changes in the hippocampus, we next investigated whether *Adnp* deficiency was accompanied by transcriptional alterations. We performed bulk RNA sequencing of hippocampal tissue from male and female WT and HET mice treated with DD vehicle, analysing each sex separately to define genotype-associated transcriptional signatures.

In male mice, 120 differentially expressed genes (FDR < 0.05) were identified between WT and HET littermates (additional file S14). Hierarchical clustering of the differentially expressed genes separated the two genotypes, indicating a distinct transcriptional profile associated with *Adnp* deficiency (figure 6A). Representative genotype-associated transcripts included *Col6a4* (Wald test, padj. = 0.0145; log_2_FC = +1.91), *Card10* (Wald test, padj. = 0.0102; log_2_FC = +0.88), *Phlda3* (Wald test, padj. = 0.0012; log_2_FC = +0.67), *N4bp3* (Wald test, padj. = 0.0001; log_2_FC = +0.59), *Efr* (Wald test, padj. = 0.0001; log_2_FC = +0.45), *Abtb1* (Wald test, padj. = 0.0001; log_2_FC = +0.63), *Arf6* (Wald test, padj. = 0.0016; log_2_FC = +0.45), *Fam117a* (Wald test, padj. = 1.28 x 10^-10^; log_2_FC = +10.12), *Rec8* (Wald test, padj. = 0.0001; log2FC = +1.44), and *mt-Nd4* (Wald test, padj. = 0.0053; log2FC = -0.61). Functional enrichment analysis revealed prominent alterations in mitochondrial and energy metabolism-related processes, including mitochondrial ATP synthesis (padj. = 0.0008; NES = -1.86; set size = 73), NADH dehydrogenase complex assembly (padj. = 0.0009; NES = -1.87; set size = 61), mitochondrial respiratory chain complex assembly (padj. = 0.0009; NES = -1.87; set size = 61), aerobic respiration (padj. = 3.10 x 10^-5^; NES = -1.81; set size = 182), oxidative phosphorylation (padj. = 5.81 x 10^-5^; NES = -1.93; set size = 141), and electron transport chain activity (padj. = 5.49 x 10^-5^; NES = -1.91; set size = 116), among others (figure 6B).

**Figure 6.**
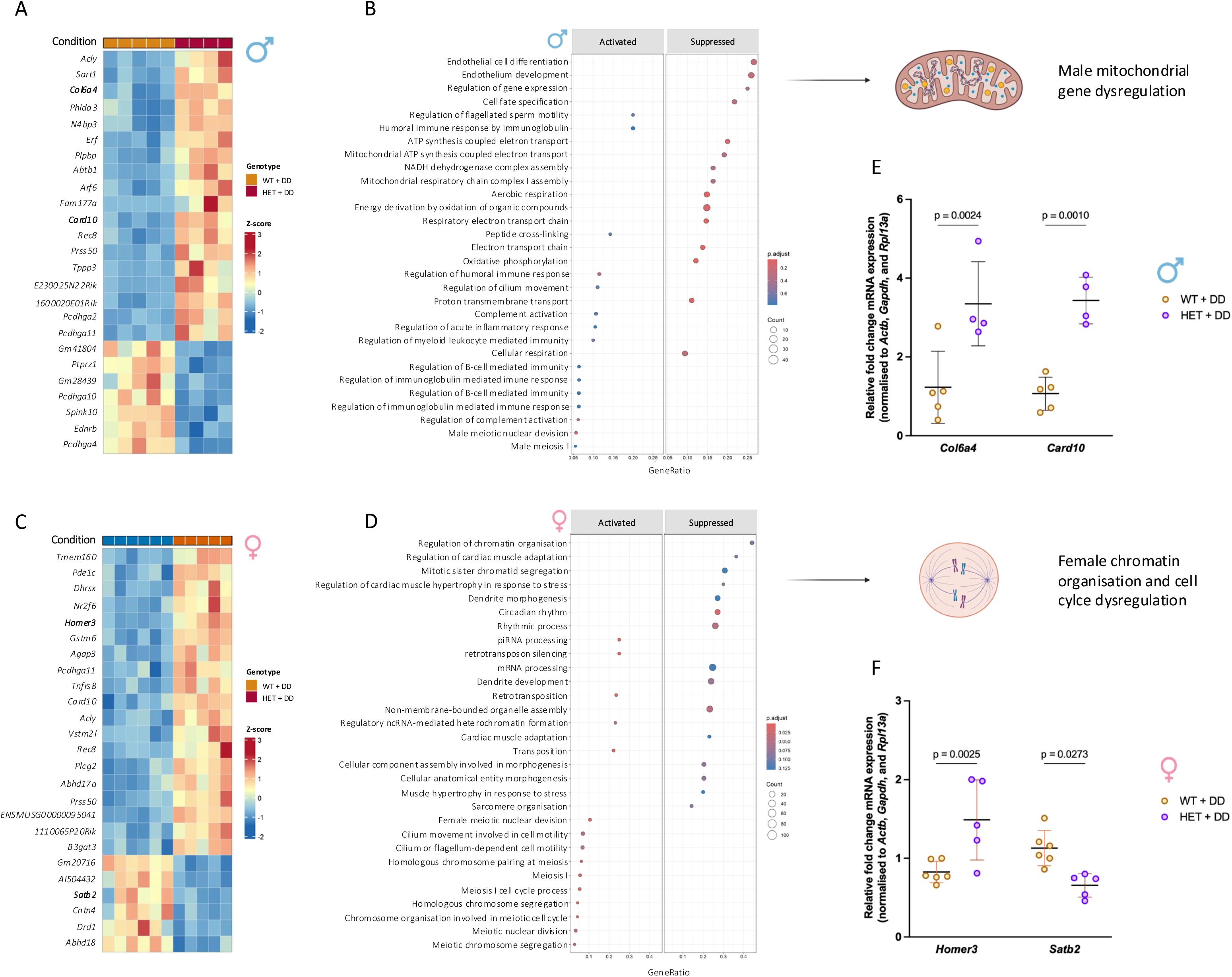
Sex-specific transcriptional alterations in the hippocampus of *Adnp*-deficient mice. **(A)** Heatmap showing representative genotype-associated transcripts in male HET mice compared with male WT littermates following DD treatment. Samples were hierarchically clustered according to their expression profiles. Genotype and treatment are indicated above the heatmap. **(B)** Functional enrichment analysis of DEGs in male HET mice, highlighting alterations in mitochondrial respiration, oxidative phosphorylation, electron transport chain activity, and energy metabolism. **(C)** Heatmap showing representative genotype-associated transcripts in female HET mice compared with female WT littermates following DD treatment. Samples were hierarchically clustered according to their expression profiles. **(D)** Functional enrichment analysis of DEGs in female HET mice, highlighting processes related to chromatin organisation, RNA processing, cell-cycle regulation, and meiotic chromosome organisation. **(E,F)** RT-qPCR validation of representative genotype-associated transcriptional changes in males **(E)** and female **(F)** mice. *Col6a4* and *Card10* expression was significantly altered in male HET mice compared with male WT littermates, whereas *Homer3* and *Satb2* expression was significantly altered in female HET mice compared with female WT littermates. RNA-sequencing analyses were performed separately in male and female mice (FDR < 0.05). RT-qPCR data are presented as mean ± s.d. and were analysed using a two-way ANOVA test with Šídák’s multiple comparisons test. (male WT + DD, n = 5; male HET + DD, n = 4; female WT + DD, n = 6; female HET + DD, n = 5).

In female mice, 245 DEGs (FDR < 0.05) were identified between WT and HET littermates (additional file S15). Hierarchical clustering similarly demonstrated genotype-associated differences in the hippocampal transcriptome (figure 6C). Representative genotype-associated transcripts included *Homer3* (Wald test, padj. = 0.0021; log_2_FC = +0.77), *Gstm6* (Wald test, padj. = 1.57 x 10^-8^; log_2_FC = +0.96), *Agap3* (Wald test, padj. = 8.39 x 10^-^5; log_2_FC = +0.31), *Pcdhga11* (Wald test, padj. = 0.0002; log_2_FC = +0.64), *Card10* (Wald test, padj. = 3.59 x 10^-5^; log_2_FC = +0.94), *Rec8* (Wald test, padj. = 8.37 x 10^-6^; log_2_FC = +1.66), *Satb2* (Wald test, padj. = 0.0011; log_2_FC = -0.86), *Cntn4* (Wald test, padj. = 0.0002; log_2_FC = -0.72), *Drd1* (Wald test, padj. = 0.0001; log_2_FC = -1.07), and *Abhd18* (Wald test, padj. = 3.25 x 10^-5^; log_2_FC = -1.13). In contrast to the male transcriptional profile, pathway enrichment in females was dominated by processes related to chromatin organisation (padj. = 0.0004; NES = -1.89; set size = 34), RNA processing (padj. = 0.0006; NES = -1.43; set size = 458), cell-cycle regulation (padj. = 7.68 x 10^-6^; NES = +2.13; set size = 18), and meiotic chromosome organisation (padj. = 4.85 x 10^-5^; NES = +2.07; set size = 69), among others (figure 6D). Thus, although *Adnp* deficiency altered hippocampal transcription in both sexes, the predominant biological processes affected differed between males and females.

We subsequently validated representative genotype-associated transcriptional changes by RT-qPCR. In males, *Col6a4* expression (male WT + DD: 1.23 ± 0.92; male HET + DD: 3.35 ± 1.07; 95% CI: -3.43 to -0.81; two-way ANOVA, p = 0.0024) and *Card10* expression (male WT + DD: 1.07 ± 0.42; male HET + DD: 3.41 ± 0.59; 95% CI: -3.68 to -1.05; two-way ANOVA, p = 0.0010) was significantly altered in HET mice compared with WT littermates (figure 6E). In females, *Homer3* expression (female WT + DD: 0.83 ± 0.14; female HET + DD: 1.49 ± 0.51; 95% CI: -1.08 to -0.24; two-way ANOVA, p = 0.0025) and *Satb2* expression (female WT + DD: 1.13 ± 0.22; female HET + DD: 0.66 ± 0.15; 95% CI: 0.05 to 0.89; two-way ANOVA, p = 0.0273) similarly showed genotype-associated changes (figure 6F).

To assess convergence between the sex-specific transcriptional responses, we compared the DEG sets. Only 13 DEGs were shared between males and females: *Acly*, *Arf6*, *Card10*, *Dnah11*, *E230025N22Rik*, *Gm20716*, *Gm42281*, *Grhpr*, *Pcdhga11*, *Pcdhga2*, *Pcdhga6*, *Prss50*, and *Rec8*. These represented 10.8% of male and 5.3% of female DEGs (Jaccard index = 3.7%). Despite this limited absolute overlap, the shared gene set was significantly greater than expected by chance (7.61-fold enrichment; odds ratio = 8.86, 95% CI: 4.90 to 15.99; Fisher’s exact test, p = 1.6 × 10^-8^). Thus, *Adnp* deficiency was predominantly associated with sex-specific transcriptional responses, with a small but statistically non-random core of shared genes. This limited transcriptional convergence contrasted with the substantially greater overlap observed at the DNA methylation level, suggesting greater sex-specific divergence at the transcriptional level than at the methylation level.

We next investigated whether NAP treatment modified that hippocampal transcriptional response associated with *Adnp* deficiency. Bulk RNA sequencing was performed on hippocampal tissue from NAP-treated WT and HET mice and compared with DD vehicle-treated animals of the same sex. A linear mixed model incorporating genotype, treatment, and the genotype × treatment interaction was used to identify genes showing a genotype-dependent response to NAP.

In male mice, 119 genes showed a significant genotype x interaction term (FDR < 0.05), of which 56 were upregulated and 63 downregulated, indicating that the transcriptional response to NAP treatment differed between male WT and HET mice (additional file S16). Hierarchical clustering of these genes demonstrated partially distinct expression patterns across genotype and treatment groups (additional figure S17A). Representative transcripts showing a significant genotype-dependent response to NAP included *Arf6* (LRT, padj. = 5.42 × 10⁻⁶; log2FC = -0.75), *1600020E01Rik* (LRT, padj. = 8.73 × 10⁻⁶; log2FC = -1.52), *Incenp* (LRT, padj. = 7.18 × 10⁻⁵; log2FC = -1.59), *Sart1* (LRT, padj. = 1.03 × 10⁻⁴; log2FC = -0.62), *Erf* (LRT, padj. = 1.03 × 10⁻⁴; log2FC = -0.80), *Gabra2* (LRT, padj. = 1.87 × 10⁻⁴; log2FC = +0.80), *Phlda3* (LRT, padj. = 3.03 × 10⁻⁴; log2FC = -1.16), *Paqr4* (LRT, padj. = 9.34 × 10⁻⁴; log2FC = -0.90), *Cpne9* (LRT, padj. = 0.0011; log2FC = -1.92), and *Map2* (LRT, padj. = 0.0259; log2FC = +0.51). Functional enrichment analysis of genes showing a genotype-dependent response to NAP revealed suppression of pathways related to cell-cycle progression and mitotic regulation, including APC/C-mediated degradation of cell-cycle proteins (padj. = 0.038; NES = -1.88; set size = 62), as well as mRNA splicing pathways (padj. = 0.0252; NES = -1.73; set size = 177). In contrast, pathways related to neuronal signalling were activated, including neurotransmitter receptor and postsynaptic density signalling (padj. = 0.0254; NES = +1.39; set size = 106), transmission across chemical synapses (padj. = 0.0262; NES = +1.38; set size = 160), trafficking of GluR2-containing AMPA receptor (padj. = 0.0454; NES = +1.52; set size = 15), and RHO GTPase cycling (padj. = 0.00072; NES = +1.30; set size = 306) (additional figure S17B). We subsequently validated *Map2* expression by RT-qPCR. *Map2* mRNA levels did not differ between male WT and HET mice under DD-vehicle treatment (male WT + DD: 1.00 ± 0.11; male HET + DD: 0.80 ± 0.24; 95% CI: -0.17 to 0.58; two-way ANOVA, p = 0.4024). Following NAP treatment, *Map2* expression was significantly higher in HET than WT mice (male WT + NAP: 0.67 ± 0.16; male HET + NAP: 1.44 ± 0.25; 95% CI: -1.14 to -0.40; two-way ANOVA, p = 0.0002). Within HET mice, NAP treatment also significantly increased *Map2* expression compared with DD-vehicle treatment (male HET + DD: 0.80 ± 0.24; male HET + NAP: 1.44 ± 0.25; 95% CI: -1.03 to -0.25; two-way ANOVA, p = 0.0016), (additional figure S17C).

In female mice, only four genes showed a significant genotype × treatment interaction (FDR < 0.05): *Baz1a* (LRT, padj. = 0.0473; log2FC = +0.02), *Tyw3* (LRT, padj. = 0.0033; log2FC = -1.34), *Gm28438* (LRT, padj. = 1.05 x 10^-^11; log2FC = -0.002), and *Gm56350* (LRT, padj. = 0.0033; log2FC = -0.0004) (additional file S18). Of these, *Tyw3* showed a substantial transcriptional effect, whereas the remaining genes exhibited minimal changes in expression despite statistical significance. The markedly smaller number of interaction-associated genes than, together with the limited magnitude of these transcriptional changes, indicates an attenuated genotype-dependent transcriptional response to NAP in female *Adnp* frameshift mice. Together, these findings indicate that NAP treatment elicited a substantially more pronounced genotype-dependent transcriptional response in male than in female *Adnp* frameshift mice.

Given the sex-specific transcriptional responses observed in female HET mice, we next asked whether oestrous-cycle-associated hormonal state might contribute to these differences. We examined hippocampal expression of the oestrogen receptor *Esr1*, progesterone receptor *Pgr*, and the oestrogen-responsive neurotrophic factor *Bdnf* in the same RNA-seq cohort, using male mice as a non-cycling reference (additional file S19). Female *Esr1* and *Pgr* expression did not differ from the corresponding male groups across genotype and treatment conditions (LRT, padj. > 0.80). *Bdnf* expression also showed no consistent genotype- or treatment-associated pattern (LRT, padj. > 0.05). Overall, the combined expression profile did not indicate a pronounced high-oestrogen transcriptional state in the female cohort at the time of tissue collection. These findings provide supportive, but not definitive, evidence that the female-specific transcriptional phenotype was not associated with a marked difference in oestrogenic state, at least for the tested genes.

### Reproductive tissue transcriptional alterations associated with Adnp deficiency

Given the molecular alterations identified in the hippocampus, we next investigated whether *Adnp* expression itself was influenced by sex or gonadal hormonal state and whether *Adnp* deficiency was associated with transcriptional alterations in reproductive tissues. We first analysed publicly available hippocampal transcriptomic data from the Four Core Genotypes (4CG) mouse model, which genetically separates sex chromosome complement from gonadal sex through the presence or absence of the *Sry* transgene (32). XX and XY-mice develop ovaries, whereas XXSry and XY-Sry mice develop testes, enabling the independent contribution of sex chromosome complement and gonadal hormonal state to be examined (figure 7A,B). Differential expression analysis in the hippocampus of 12-month-old mice did not identify *Adnp* as significantly regulated by either sex chromosome complement or gonadal state (figure 7C). Consistent with this, normalised *Adnp* expression did not differ between XY- and XX mice with ovaries (XY-: 81.47 ± 30.33; XX: 100.24 ± 38.45; 95% CI: -64.43 to 26.89; LRT, padj. = 0.6917) or between XY-Sry and XXSry mice with testes (XY-Sry: 97.96 ± 51.84; XXSry: 114.23 ± 33.92; 95% CI: -59.81 to 27.27; LRT, padj. = 0.7504). Similarly, no differences were observed between ovarian and testicular states within either the XY (XY-: 81.47 ± 30.33; XY-Sry: 97.96 ± 51.84; 95% CI: -62.15 to 29.17; LRT, padj. = 0.7693) or XX (XX: 100.24 ± 38.45; XXSry: 114.23 ± 33.92; 95% CI: -57.53 to 29.55; LRT, padj. = 0.8254) chromosome complement (figure 7D). We next asked whether *Adnp* might nevertheless contribute to sex- or hormone-associated transcriptional programmes through its reported transcription factor activity (35). Transcription factor enrichment analysis using ChEA3 identified *Adnp* as a candidate regulator of a subset of sex- and hormone-associated genes in the 4CG hippocampus, with 11 of 82 DEGs overlapping the predicted *Adnp* regulon (rank 219 of 1,632 transcription factors queried). However, the enrichment was substantially weaker than that of the highest-ranking regulators (additional file S20). These findings suggest that *Adnp* expression itself is not detectably regulated by sex chromosome complement or gonadal hormonal state in the hippocampus of 12-month-old animals. Although the 4CG analysis did not account for the stage of the oestrous cycle, which could contribute to within-sex transcriptional variability, the findings are consistent with a possible, modest contribution of *Adnp* to downstream hormone-associated transcriptional programmes.

**Figure 7.**
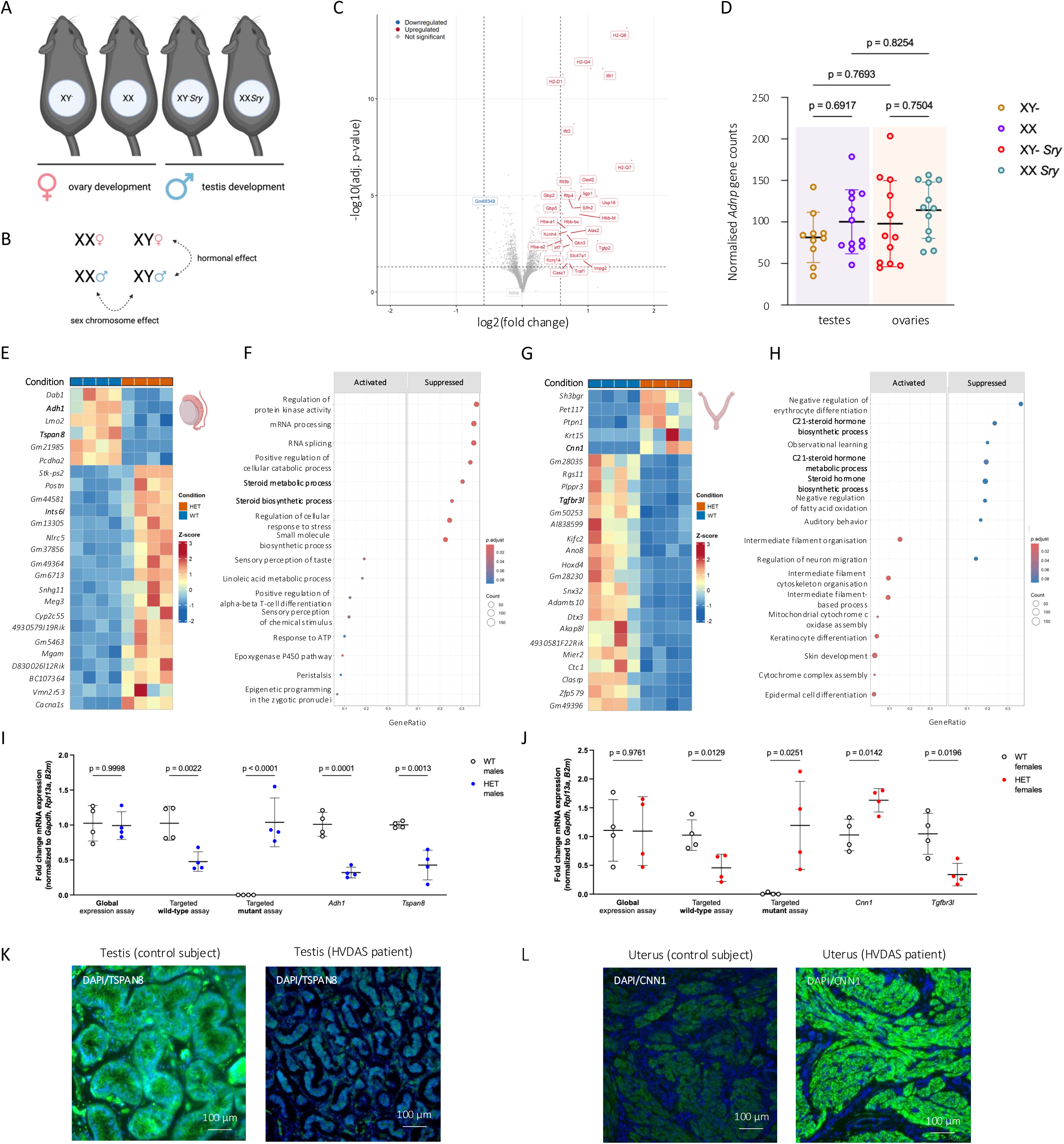
Reproductive tissue transcriptional alterations associated with *Adnp* deficiency. **(A,B)** Schematic of the Four Core Genotypes (4CG) model and its application to distinguish sex chromosome and gonadal effects on hippocampal *Adnp* expression. XX and XY-Sry mice develop ovaries, whereas XXSry and XY-Sry mice develop testes, enabling assessment of sex chromosome complement independently of gonadal sex. **(C)** Differential expression analysis of hippocampal transcriptomes from 4CG mice showing sex- and gonadal sex-associated transcriptional changes. *Adnp* was not significantly differentially expressed. **(D)** Normalised *Adnp* expression across 4CG groups. No significant differences were detected between sex chromosome complements or gonadal states (LMM, FDR < 0.05). **(E)** Heatmap showing representative genotype-associated DEGs in testis from HET mice compared with WT littermates. Samples were hierarchically clustered according to their expression profiles. **(F)** Functional enrichment analysis of DEGs in testis, highlighting alterations in cellular regulation, RNA processing and reproductive processes, including steroid hormone biosynthetic processes. **(G)** Heatmap showing representative genotype-associated DEGs in uterus from HET mice compared with WT littermates. **(H)** Functional enrichment analysis of DEGs in uterus, highlighting alterations in epithelial differentiation, extracellular matrix organisation, cell adhesion and steroid hormone biosynthesis. **(I,J)** Targeted RT-qPCR validation of representative genotype-associated transcriptional changes in testis **(I)** and uterus **(J)**. *Adh1* and *Tspan8* were selected as representative testicular transcripts, whereas *Cnn1* and *Tgfbr3l* were selected as representative uterine transcripts based on robust genotype-associated changes identified by RNA sequencing and their relevance to tissue-specific cellular and reproductive processes. mRNA expression was normalised to *Gapdh*, *Rpl13a,* and *B2m*. Statistical analysis was performed using Welch’s t-test. Data are presented as mean ± s.d. (n = 4 per genotype and tissue). **(K,L)** Representative immunohistochemical staining of TSPAN8 in human testicular tissue **(K)** and CNN1 in human uterine tissue **(L)** from control subjects and individuals with Helsmoortel-Van der Aa syndrome. Scale bars, 100 μm.

We therefore investigated whether the heterozygous *Adnp* frameshift variant alters transcriptional programmes in reproductive tissues, with particular focus on pathways that could provide a common molecular link between sex-dependent phenotypes identified in the hippocampus. Bulk RNA sequencing was performed separately in testis and uterus from WT and HET mice, consistent with the sex-stratified approach used for hippocampal transcriptomic analyses. In testis, *Adnp* deficiency produced a distinct transcriptional profile, with genotype-associated separation observed by hierarchical clustering (figure 7E). Functional enrichment analysis identified alterations in cellular protein kinase activity (padj. = 0.0047; NES = -1.79; set size = 354), mRNA processing (padj. = 0.0063; NES = -1.67; set size = 477), and reproductive processes, including steroid hormone biosynthetic processes (padj. = 0.0047; NES = -1.95; set size = 142) (figure 7F). A distinct genotype-associated transcriptional profile was likewise observed in uterine tissue (figure 7G), with enriched processes including suppression of steroid hormone biosynthesis processes (padj. = 0.0007; NES = -1.83; set size = 31), among others. In contrast, pathways related to intermediate filament organisation (padj. = 0.0001; NES = +2.21; set size = 33), cytoskeleton organisation (padj. = 0.0002; NES = +2.21; set size = 53) and skin development (padj. = 0.0003; NES = +1.91; set size = 259) were activated (figure 7H). Notably, despite the distinct tissue contexts of testis and uterus, steroid hormone biosynthesis was enriched in both reproductive tissues, identifying steroid hormone metabolism as a shared transcriptional feature of *Adnp* frameshift mice. Additional concordant alterations were observed in RNA splicing (uterus, padj. = 0.024; testis, padj. = 0.0063) and motile cilium assembly (uterus, padj. = 0.0171; testis, padj. = 0.0447), both of which were downregulated in HET mice. Cytoplasmic translational initiation was also altered in both tissues (uterus, padj. = 0.0049; testis, padj. = 0.0013), although its direction differed between tissues, being increased in uterus and decreased in testis. Thus, while the broader transcriptional responses remained tissue-specific, several biological processes were shared across reproductive tissues.

To validate representative RNA-sequencing findings, we selected transcripts implicated in cellular differentiation and reproductive tissue function. In testis, *Adh1* (WT: 1.01 ± 0.17; HET: 0.32 ± 0.08; 95% CI: -0.95 to -0.43; Welch’s t-test, p = 0.0001) and *Tspan8* (WT: 1.00 ± 0.05; HET: 0.43 ± 0.21; 95% CI: -0.90 to -0.25; Welch’s t-test, p = 0.0013) were significantly reduced in HET mice compared with WT littermates (figure 7I). In uterus, *Cnn1* (WT: 1.03 ± 0.27; HET: 1.63 ± 0.20; 95% CI: 0.18 to 1.03; Welch’s t-test, p = 0.0142) was increased, whereas *Tgfbr3l* (WT: 1.05 ± 0.18; HET: 0.34 ± 0.10; 95% CI: -1.24 to -0.17; Welch’s t-test, p = 0.0196) was reduced in HET mice (figure 7J). We next examined *Adnp* transcript abundance directly in reproductive tissues using complementary assays designed to distinguish total *Adnp* expression from transcripts retaining or spanning the deleted 14-bp region (21). Total *Adnp* transcript abundance was unchanged in both testis (WT: 1.03 ± 0.25; HET: 0.99 ± 0.20; 95% CI: -0.43 to 0.37; Welch’s t-test, p = 0.9998) and uterus (WT: 1.11 ± 0.53; HET: 1.10 ± 0.60; 95% CI: -1.00 to 0.97; Welch’s t-test, p = 0.9761). In contrast, transcripts detected by the wild-type-specific assay were significantly reduced in both testis (WT: 1.03 ± 0.24; HET: 0.48 ± 0.14; 95% CI: -0.91 to -0.18; Welch’s t-test, p = 0.0022) and uterus (WT: 1.03 ± 0.27; HET: 0.46 ± 0.24; 95% CI: -1.00 to -0.13; Welch’s t-test, p = 0.0129), whereas mutant transcripts were detected exclusively in HET tissues (testis WT: 0.00 ± 0.00; testis HET: 1.04 ± 0.35; 95% CI: 0.48 to 1.59; Welch’s t-test, p < 0.0001; uterus WT: 0.01 ± 0.02; uterus HET: 1.31 ± 0.63; 95% CI: 0.31 to 2.31; Welch’s t-test, p = 0.0251).

To assess whether selected transcriptional alterations identified in mice were also observed in human reproductive tissues, we examined TSPAN8 and CNN1 protein expression in testicular and uterine tissues from individuals with Helsmoortel-Van der Aa syndrome and control subjects. Immunohistochemical analysis showed reduced TSPAN8 expression in testicular tissue from the analysed individual with Helsmoortel-Van der Aa syndrome compared with controls (CTR: 1.00 ± 0.19; HVDAS: 0.50 ± 0.01; 95% CI: -0.96 to -0.04; Welch’s t-test, p = 0.0432), consistent with the reduced *Tspan8* expression observed in HET mouse testis (figure 7K). Conversely, CNN1 expression was increased in uterine tissue from the analysed individual with Helsmoortel-Van der Aa syndrome compared with controls (CTR: 1.00 ± 0.22; HVDAS: 3.78 ± 0.02; 95% CI: 2.25 to 3.32; Welch’s t-test, p = 0.0019), concordant with increased *Cnn1* expression observed in HET mouse uterus (figure 7L). Targeted transcript analyses further identified increased *ADH1C* expression in human testicular tissue (CTR: 1.51 ± 1.63; HVDAS: 3.51 ± 0.77; 95% CI: -3.69 to -0.32; Welch’s t-test, p = 0.0189) and increased *TGFBR3L* expression in human uterine tissue (CTR: 1.00 ± 0.11; HVDAS: 1.75 ± 0.56; 95% CI: -1.34 to -0.15; Welch’s t-test, p = 0.0135), representing changes in the opposite direction to those observed in the corresponding mouse tissues (additional file S21).

Collectively, these findings extend the molecular consequences of *Adnp* deficiency beyond the hippocampus to reproductive tissues. *Adnp* expression itself was not detectably regulated by sex chromosome complement or gonadal hormonal state in the 4CG mouse hippocampus, although *Adnp* emerged as a modest candidate regulator of a subset of hormone-associated transcriptional changes. In contrast, *Adnp* deficiency was associated with distinct transcriptional alterations in testis and uterus, with convergence on steroid hormone biosynthesis across both tissues. Together with the sex-dependent molecular alterations identified in the hippocampus, these findings implicate altered steroid hormone-related biology as a potential shared downstream component of *Adnp* deficiency, while emphasising that the specific transcriptional response remains strongly dependent on tissue context.

### No sex-dependent clinical features in individuals with Helsmoortel-Van der Aa syndrome

To determine whether the molecular differences associated with *Adnp* deficiency in male and female mice were reflected in the clinical phenotype of Helsmoortel-Van der Aa syndrome, we re-analysed clinical data from a cohort of 129 individuals with genetically confirmed *ADNP* variants (11), comprising 69 males and 60 females. Across the broad range of neurological, developmental, behavioural, endocrinological, and systemic features assessed, clinical manifestations were largely shared between sexes, with no feature showing a significant sex-associated difference after correcting for multiple testing (Fisher’s exact test, FDR-adj. q > 0.05 for all features) (figure 8A). Similarly, developmental measures showed no statistical sex-dependent differences after correction for multiple testing, including age at first words (males: 26.22 ± 12.60 months; females: 36.25 ± 16.77 months; 95% CI: 1.99 to 18.07; Student’s t-test, p = 0.0123, q = 0.1575) and age at independent walking (males: 31.41 ± 14.22 months; females: 27.35 ± 9.79 months; 95% CI: -9.48 to 1.35; Student’s t-test, p = 0.3162, q = 0.4795) (figure 8B). Overall, no robust sex-dependent differences were identified across the clinical or developmental features assessed.

**Figure 8.**
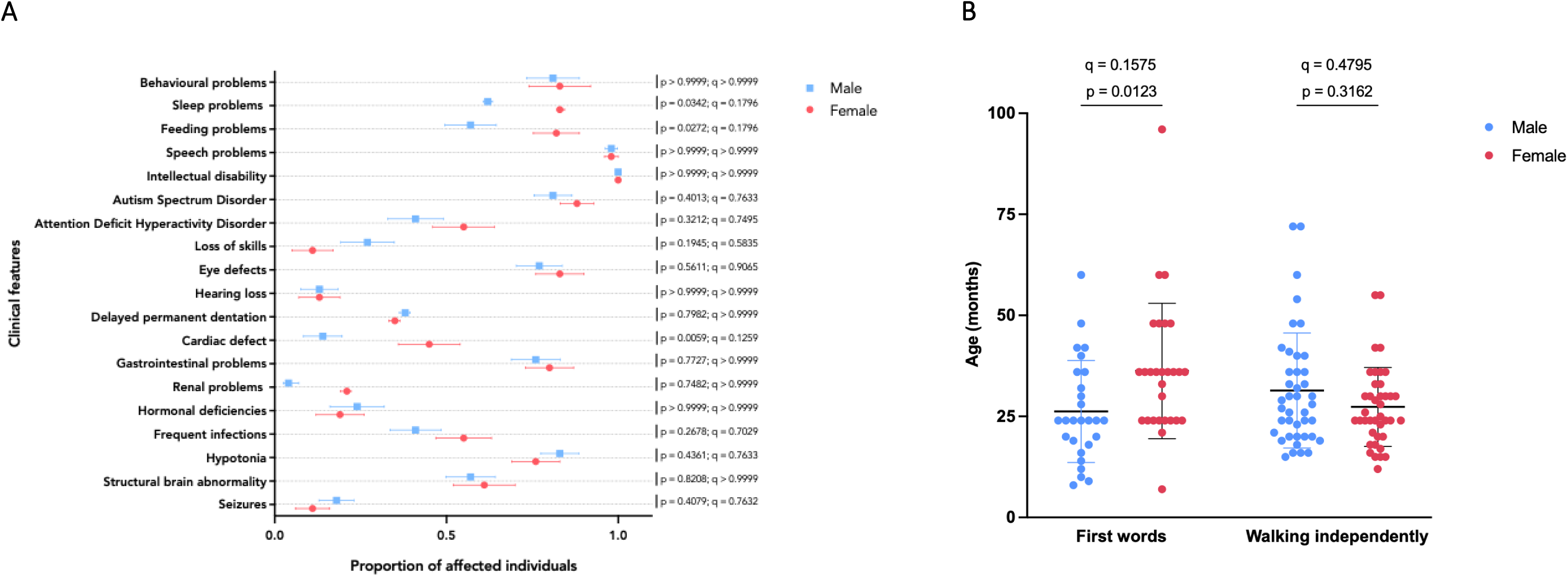
Sex-stratified clinical and developmental features in individuals with Helsmoortel-Van der Aa syndrome. Clinical and developmental characteristics were assessed in 129 individuals with genetically confirmed *ADNP* variants, comprising 69 males and 60 females. **(A)** Proportion of male and female individuals presenting with selected neurological, developmental, behavioural, endocrinological, dysmorphic, and systemic clinical features. Categorical variables were analysed using Fisher’s exact test, with false discovery rate correction for multiple testing (FDR < 0.05). Data are presented as the proportion of affected individuals, with 95% CIs. No clinical feature showed a significant sex-associated difference after FDR correction (all q > 0.05). **(B)** Age at first words and age at independent walking in male and female individuals. Continuous variables were analysed using Student’s t-test, followed by FDR correction across the developmental measures assessed. Data are presented as mean ± s.d. Age at first words showed a nominal sex-associated difference (p = 0.0123), which did not remain significant after FDR correction (q = 0.1575), whereas age at independent walking did not differ between sexes (p = 0.3162; q = 0.4795).

## Discussion

In this study, we investigated how sex influences the behavioural and molecular consequences of *Adnp* deficiency, whether these effects are modified by NAP treatment, and whether these findings are reflected in individuals with Helsmoortel-Van der Aa syndrome. *Adnp* deficiency produced a broadly shared behavioural phenotype in male and female mice, but the features contributing to this phenotype differed between sexes. At the molecular level, genotype was the predominant determinant of hippocampal DNA methylation, whereas transcriptional responses showed greater sex-dependent divergence, including distinct enrichment of mitochondrial pathways in males and chromatin, RNA-processing and cell-cycle-related pathways in females. NAP partially normalised behavioural organisation in both sexes and increased hippocampal ADNP protein abundance in HET mice but did not measurably reverse the underlying methylation landscape and produced only limited transcriptional changes. In reproductive tissues, *Adnp* deficiency was associated with convergent alterations in steroid hormone biosynthetic pathways in testis and uterus, despite no evidence that hippocampal *Adnp* expression was directly regulated by sex chromosome complement or gonadal hormonal state. Selected molecular alterations were also observed in human reproductive tissues. Finally, re-analysis of the Helsmoortel-Van der Aa syndrome clinical cohort identified no robust sex-associated differences across the clinical and developmental features examined after correction for multiple testing. Thus, at the global clinical level, the predominantly shared phenotype in individuals with Helsmoortel-Van der Aa syndrome contrasts with the sex-dependent behavioural organisation observed in mice. However, detailed neuropsychological and behavioural phenotyping was not available in the human cohort, precluding direct comparison of the finer-scale behavioural features identified in mice. Collectively, these findings indicate that sex influences the molecular and behavioural consequences of *Adnp* deficiency without being sufficient to define a distinct clinical presentation based on the features assessed here.

### Sex modifies the behavioural organisation of *Adnp* deficiency

Male and female frameshift mice showed a comparable overall displacement of behavioural organisation from their respective WT controls, indicating that *Adnp* deficiency produces a shared behavioural phenotype across sexes. However, the individual features contributing to this phenotype differed. Male HET mice showed greater alterations in exploratory activity and physical social contact, whereas females showed more pronounced changes in social approach, withdrawal and group organisation. Thus, sex influenced the organisation of the phenotype rather than its overall presence or magnitude.

These findings are consistent with the heterogeneous behavioural phenotypes reported across *Adnp* murine models. Sex-dependent differences in cognition, gait, social behaviour and peripheral molecular phenotypes have been described in haploinsufficient (Δexons 1-6) (14), p.Tyr718* mutant mice (15), and *Adnp* frameshift mice (45), whereas another *Adnp* haploinsufficient (Δexon 6) model has shown broadly similar behavioural impairments between males and females (17). Differences between models may reflect the underlying *Adnp* variant (46,47), residual protein function (36), developmental consequences (15), age (12,14), the genetic background of the mouse model (48), and interactions with laboratory environment and experimental design (49). The present findings extend this literature by showing that sex-dependent behavioural organisation can occur even when the overall magnitude of genotype-associated behavioural displacement is similar between males and females.

The biological plausibility of this sex-dependent organisation is supported by evidence that ADNP expression and function vary according to sex and tissue context. Sex- and oestrous-cycle-associated differences in *Adnp* expression have been reported in the hypothalamus (18), while higher *ADNP* expression has been described in male compared with female hippocampus in both mice and humans (13). However, our Four Core Genotypes analysis did not support direct regulation of hippocampal *Adnp* expression by gonadal hormonal state or sex chromosome complement. These findings suggest that sex-associated differences in ADNP biology may instead arise through tissue-specific regulatory networks, including upstream transcriptional regulators, rather than through direct control of *Adnp* itself. ADNP has also been implicated in sex-dependent neuronal morphogenesis, cortical connectivity and calcium signalling (50), as well as sex-dependent and variant specific circadian rhythmicity (51) and age- and tissue-specific regulation of sex chromosome genes (52). Together, these observations support a model in which sex modifies the cellular context in which reduced *Adnp* function is expressed.

### *Adnp* deficiency engages shared epigenetic and divergent transcriptional programmes

Our molecular analyses distinguish the relative convergence of the methylomic response from the greater divergence of transcriptional consequences. Genotype was the predominant determinant of hippocampal DNA methylation, with extensive genotype-associated methylation changes shared between males and females. By contrast, only 13 DEGs were shared between sexes, although this overlap was greater than expected by chance. These findings suggest that *Adnp* deficiency establishes a relatively conserved epigenomic state, whereas downstream transcriptional consequences are more strongly shaped by sex, cellular context, disease-causing variants, and age (32,53,54).

Several sex-associated molecular findings reinforce this distinction. Sex-associated methylation changes involving *Nlgn1* and *Foxp2* intersect with previous evidence of these neuronal programmes, including increased *Nlgn1* expression in female *Adnp* haploinsufficient (Δexons 1-6) mice (13), and sex-dependent *Foxp2* transcript regulation (44). The present findings therefore extend observations from individual loci to a broader molecular framework in which sex-dependent epigenetic and transcriptional context may contribute to the different behavioural manifestations of *Adnp* deficiency.

The transcriptomic signatures further suggest that *Adnp* deficiency affects multiple interconnected cellular programmes rather than a single canonical pathway. Male hippocampi showed prominent alterations in mitochondrial respiration and oxidative phosphorylation, whereas female hippocampi showed greater enrichment of chromatin organisation, RNA processing and cell-cycle-associated processes. These findings are consistent with emerging evidence for sex-dependent effects of *Adnp* deficiency on histone modifications (7,55) and hippocampal neurobiology, including sex-dependent alterations in neurogenesis and cellular stress responses (16). They also provide a molecular context for the mitochondrial abnormalities previously identified in human Helsmoortel-Van der Aa autopsy tissue (35). Therefore, mitochondrial and chromatin-related pathways emerge as candidate modifiers of the sex-dependent consequences of *Adnp* deficiency, although their precise cellular and functional relevance remains to be established.

### NAP partially improves functional phenotypes without resetting the underlying molecular state

NAP (NAPVSIPQ), clinically formulated as davunetide, is the shortest biologically active peptide derived from ADNP and was originally characterised as a neuroprotective microtubule-associated peptide that enhances microtubule dynamics in neurons and astrocytes (56,57). Previous studies in *Adnp*-deficient models have reported partial rescue of developmental, synaptic and behavioural abnormalities following NAP administration (14,15). Our findings extend these observations by showing that chronic NAP treatment partially shifted the behavioural organisation of HET mice towards the WT state in both sexes, although the behavioural domains showing the greatest treatment-associated changes differed between males and females.

NAP also increased hippocampal ADNP protein abundance in HET mice of both sexes. Because mutant *Adnp* transcript abundance was not increased by treatment, this response is unlikely to reflect simple induction of the mutant allele. Rather, NAP might enhance the stability, availability, or downstream utilisation of residual functional ADNP protein, potentially through interactions with EB1/3 proteins (58), although the precise nuclear mechanism remains unresolved (59).

Importantly, the behavioural response occurred without a corresponding reversal of the widespread hippocampal methylation phenotype. NAP produced only limited transcriptional changes and did not induce a broad transcriptome-wide correction in HET hippocampi. These findings indicate that the behavioural improvement can occur without restoration of the molecular state associated with *Adnp* deficiency. Instead, NAP may engage downstream cellular processes that improve functional resilience despite persistence of the underlying epigenomic alteration, consistent with its reported effects on microtubule integrity, cellular stress responses and mitochondrial protection (60–62). This interpretation is particularly relevant to the mitochondrial transcriptional phenotype observed in male frameshift mouse hippocampus. *ADNP* deficiency has been associated with impaired mitochondrial function in human tissues (35), while NAP has been shown to protect mitochondria from stress-associated cytochrome c release (63). Recent work in haploinsufficient (Δexons 1-6) and p.Tyr718* mutant *Adnp* mice similarly identified sex-dependent mitochondrial and unfolded-protein-response abnormalities that were partially moderated by NAP (16). These observations provide a potential cellular context for the treatment response, while the extent to which these pathways mediate behavioural improvement remains unresolved.

### Reproductive tissues identify a potential endocrine component of *Adnp* deficiency

The reproductive-tissue analyses provide a complementary perspective on how sex-dependent biology may intersect with *Adnp* deficiency. The Four Core Genotypes analysis did not provide evidence that hippocampal *Adnp* expression is directly regulated by sex chromosome complement or gonadal hormonal state. Although *ADNP* emerged as a candidate regulator of a subset of hormone-associated transcripts, the relatively modest enrichment argues against a dominant upstream determinant of the sex- or hormone-associated transcriptional programme.

Instead, the testis and uterine transcriptomes of HET mice pointed towards a downstream relationship. Despite their markedly different cellular composition and physiological functions, both tissues showed altered steroid hormone biosynthetic pathways following *Adnp* deficiency. Additional convergence was observed in RNA splicing and motile cilium assembly, although the broader transcriptional responses remained strongly tissue specific. These findings raise the possibility that altered steroid hormone-related biology represents a shared downstream component of *Adnp* deficiency rather than a consequence of direct hormonal regulation of *Adnp* itself. This interpretation is compatible with earlier evidence linking *Adnp* to steroid hormone-associated pathways (20). The findings therefore extend the biological consequences of *Adnp* deficiency beyond the central nervous system and identify endocrine and reproductive pathways as potential tissue-specific components of the phenotype.

### Molecular sex differences do not define the clinical syndrome

The clinical cohort analysis provides an important translational counterpoint to the sex-dependent behavioural and molecular differences observed in mice. No robust sex-associated differences were identified across the clinical and developmental features assessed in 129 individuals with Helsmoortel-Van der Aa syndrome after correction for multiple testing, consistent with the predominantly shared clinical presentation of the syndrome (6,10,11). This finding does not establish molecular equivalence between males and females (64). The mouse analysis identified sex-dependent differences in specific behavioural features for which there is currently no equivalent phenotypic resolution in the human cohort. Detailed neuropsychological and behavioural phenotyping will therefore be required to determine whether comparable sex-dependent differences are present in individuals with Helsmoortel-Van der Aa syndrome. Limited evidence from a small longitudinal cohort of 15 individuals with Helsmoortel-Van der Aa syndrome, in which the two variants represented in both sexes were compared, suggested better adaptive and communicative performance in females than males (65). Although preliminary, these findings indicate that more granular sex-associated phenotypes warrant further investigation. More broadly, biological sex can influence cellular pathways, gene regulation and treatment responses without producing a readily detectable difference in the clinical phenotype (66,67). Thus, these findings support a model in which sex-dependent molecular and behavioural effects can coexist with a largely shared clinical syndrome, potentially modifying disease biology without being primary determinants of clinical expression (68).

### Sex-dependent pharmacological responses warrant consideration in therapeutic development

The combined preclinical and clinical evidence raises the possibility that sex may modify pharmacological responses in *ADNP* deficiency. In the present study, NAP partially improved behavioural abnormalities in both male and female HET mice, but produced a markedly broader transcriptional response in males. However, this pattern should not be interpreted as evidence for generally greater treatment responsiveness in males. In the same *Adnp* frameshift mouse model, LSD1 inhibition produced robust restoration of excitatory and inhibitory synaptic transmission and normalisation of activating and repressive histone marks in females, whereas males showed more limited effects on synaptic function and synaptic gene expression (69). The contrasting direction of these responses suggests that sex may modify pharmacological sensitivity in an intervention-dependent manner rather than confer a uniform treatment advantage to either sex.

Hormonal state represents one potential contributor to such differences, although this remains untested in the present study. Intranasal NAP uptake in female mice has recently been reported to vary across the oestrous cycle (70). In our cohort, hippocampal *Esr1*, *Pgr*, and *Bdnf* expression did not indicate a pronounced high-oestrogen state, but these observations cannot substitute for direct assessment of oestrous stage or circulating hormone concentrations. Accordingly, hormonal state could plausibly contribute to differences in NAP exposure or response.

The clinical history of davunetide provides additional, albeit indirect, translational context. Davunetide has demonstrated clinical safety and signals of benefit in some human neurodegenerative settings (71). The original randomised trial in progressive supranuclear palsy, which included male and female participants, did not demonstrate overall efficacy (72), whereas subsequent sex-stratified analyses reported treatment-associated benefits in women (73). These findings arose in a different disease context and were not prospectively designed to test sex differences, and therefore cannot establish sex-dependent pharmacological efficacy of davunetide. Nevertheless, together with the contrasting sex-dependent pharmacological responses observed across *Adnp* mouse studies, they support prospective consideration of sex as a prespecified biological variable in future therapeutic studies of Helsmoortel-Van der Aa syndrome. In paediatric trials, developmental stage and potentially hormonal context may also need to be considered. Prospective studies incorporating these variables will be required to determine whether sex-dependent treatment responses observed in preclinical models translate to individuals with Helsmoortel-Van der Aa syndrome.

## Conclusion

Overall, this study identifies sex as an important modifier of the biological consequences of *Adnp* deficiency while demonstrating that the core phenotype remains fundamentally shared across sexes. Sex-dependent behavioural organisation was accompanied by divergent hippocampal transcriptional programmes, whereas the methylation response showed substantial convergence. NAP partially improved behavioural phenotypes in both sexes and increased residual wild-type ADNP protein abundance without reversing the underlying hippocampal methylation landscape. Reproductive-tissue analyses further identified steroid hormone biosynthesis as a potential shared downstream component of *Adnp* deficiency. Despite these molecular and behavioural differences, males and females with Helsmoortel-Van der Aa syndrome showed broadly overlapping clinical phenotypes in the features assessed. Together, these findings support consideration of sex and hormonal context when investigating the biological mechanisms and therapeutic responses associated with *Adnp* deficiency, while recognising that their contribution to clinical heterogeneity remains to be established.

## Limitations and caveats

Several limitations should be considered when interpreting these findings. First, the study used a single *Adnp* frameshift mouse model, and the consequences of different pathogenic *Adnp* variants may not be identical. Differences in residual protein function, transcript processing and variant position could contribute to the behavioural and molecular heterogeneity observed across published *Adnp* models. The present findings should therefore not be assumed to generalise to all genetic forms of Helsmoortel-Van der Aa syndrome. Second, reproductive-tissue analyses and human validation were constrained by sample availability. Human molecular analyses included tissue from a single individual with Helsmoortel-Van der Aa syndrome per tissue and therefore provide exploratory cross-species validation rather than evidence of population-level effects. Moreover, the relatively small numbers of animals in some transcriptomic and reproductive-tissue experiments limit statistical power for detecting modest effects and higher-order genotype × sex × treatment interactions.

Although the Four Core Genotypes analysis enabled separation of sex-chromosome complement from gonadal sex, hormonal state was not directly measured. Oestrous stage was not determined, and potential sex differences in NAP exposure or pharmacokinetics cannot therefore be excluded. For reproductive-tissue RNA-sequencing, murine uterus and testis were selected to match the tissues available from the two individuals with Helsmoortel-Van der Aa syndrome, enabling a direct cross-species comparison. These tissues are hormone sensitive but are not equivalent to the principal sites of gonadal steroid synthesis, and the testis contains a relatively small steroidogenic Leydig-cell population (74,75). The shared suppression of a hormone-biosynthesis pathway across these tissues is therefore of interest but should be interpreted cautiously given the tissue selection and limited human sample size.

Behavioural phenotyping was performed in group-housed animals using the LMT platform, and social interactions, cage composition and hierarchy may therefore influence individual behavioural measures. Cage identity was incorporated as a random effect in the relevant mixed models, but group housing remains an inherent feature of the experimental paradigm. The LMT also primarily captures spontaneous and social behaviour and does not directly assess learning and memory (22). This is relevant because sex-dependent differences in learning have been reported in the same *Adnp* mouse model using the Morris water maze (45), although NAP treatment was not examined in that context. Thus, whether the observed sex-dependent molecular effects or NAP-associated behavioural effects extend to cognitive domains remains unknown. Finally, the molecular analyses were predominantly performed in bulk tissues and therefore cannot resolve cell-type-specific responses. The human cohort analyses were also limited to clinical and developmental features and did not assess behavioural or molecular sex differences. Therefore, sex-dependent effects cannot be excluded. Accordingly, the absence of sex-associated clinical differences should not be interpreted as evidence that more subtle molecular or phenotypic sex effects are absent.

## Contributors

C.P.D. maintained the mouse colony, administered NAP treatment, and was the principal researcher responsible for performing, conceptualising, and analysing all experiments in the study. He wrote the manuscript and prepared the final figures. J.I. performed the analysis of the mouse methylation arrays and facilitated the visualisation of all related figures. M.B.V.D.L. performed RFID tag implantation and LMT recordings. L.H. performed the clinical sex-stratified re-analysis of the human Helsmoortel-Van der Aa syndrome cohort under the supervision of C.P.D., M.M., A.C.J., and E.F.; K.V.M. was the master’s student under the supervision of C.P.D. and assisted with NAP administration and participated in all experiments as part of her master’s thesis. E.E. performed mouse genotyping and conducted all gene expression assays using RT-PCR under the supervision of C.P.D.; A.Sc. is a research assistant under the supervision of C.P.D., M.M., and A.C.J. and assisted with western blotting and immunohistochemistry experiments using human reproductive tissues. A. Sch. performed the mouse methylation array as a research technician. L. Mo. and M.H. are bioinformaticians who facilitated the RNA sequencing analyses of mouse hippocampus and reproductive tissues. Z.M., M.A., and J.V. provided post-mortem tissue from the male individual with Helsmoortel-Van der Aa syndrome and clinical information. P.T. and N.G. provided histologically processed testis tissue from the male individual with Helsmoortel-Van der Aa syndrome for immunohistochemistry experiments. M.d.B.O. is a senior research technician at the Erasmus MC tissue bank and PARTS clinical trials coordinator who facilitated access to processed control tissues from testes and uterus. E.F. is a biostatistician who oversaw all statistical methods implemented in this study. L.M. facilitated the initial bioinformatic analysis of the RNA sequencing data and supervised the bioinformatics team. A.C.J. and M.M. supervised the clinical component and analyses of the project. D.J.A. facilitated the data analysis of the LMT, developed the MouseKing analytical framework, and provided all related figures. I.G. provided the experimental drug NAP (davunetide) and trained C.P.D. in NAP administration. She was involved in the collection of the male autopsy material and reviewed the manuscript. R.F.K. supervised the entire project, conceptualised all experiments with C.P.D., and reviewed the manuscript. C.P.D. and R.F.K. were involved in funding acquisition. All authors read and approved the final version of the manuscript.

## Acknowledgements

We are especially grateful to the two children and their families who contributed to this study. We express our deepest gratitude to the family who donated autopsy material following the death of their son, and to the family of the female individual who donated tissue following her hysterectomy. We thank the European ADNP Association for its continuous collaboration and support (https://adnpeurope.eu). We thank our talented master’s student, Amber Buys, for her help and assistance with NAP injections as part of the fulfilment of her master’s internship. We thank Prof. Dr. Wim Vanden Berghe for facilitating access to the pyrosequencer. We highly value the input of Prof. Dr. Jozef Gecz regarding the incorporation of the 4GC mouse model and the interpretation and integration of these results. We thank our collaborator Dr. Eliezer Giladi for sharing their expertise on NAP. During the preparation of this manuscript, generative AI and AI-assisted technologies were used to refine, edit, and format human-written text and to assist with the formatting and presentation of manuscript text. The authors reviewed and verified the resulting content and remain fully responsible for the accuracy, interpretation, and final content of the manuscript.

## Data sharing statement

The data presented in this study are available in the article and supplementary information. Raw IDAT files for genome-wide DNA methylation profiling of male and female *Adnp* mice, and raw RNA sequencing data from male and female hippocampus and reproductive tissues (murine testis and uterus), are available via Zenodo (10.5281/zenodo.15830464).

## Consent for publication

Written informed consent was obtained from the individuals included in this study, permitting clinicians and researchers from the Department of Medical Genetics, University Hospital Antwerp, to use photographs, biopsies, and/or X-rays for medical purposes, including discussion among colleagues, teaching, and publication in scientific papers.

