## Supplementary material for "The Female Side of Autism: Sexual Dichotomies impacting the Helsmoortel-Van der Aa syndrome pathology": Captions

Captions for additional files

**Additional file S1. Uncropped western blots.** Uncropped western blot images underlying the cropped blots shown in the main figures. (**A-H**): mouse hippocampus. Blots are organised by sex and biological replicate group, with ADNP (150 kDa) shown on the left (panels A-D) and the GAPDH loading control (37 kDa) shown on the right (panels E-H) for the same samples. (**A**) Males, biological replicates 1-3, ADNP. (**B**) Males, biological replicates 4-6, ADNP. (**C**) Females, biological replicates 1-3, ADNP. (**D**) Females, biological replicates 4-6, ADNP. (**E-H**) Corresponding GAPDH loading-control blots for panels A-D, respectively. Lanes in each panel are loaded WT + DD-vehicle, HET + DD-vehicle, WT + NAP, and HET + NAP, in biological triplicate per group. (**I-L**): human testis and uterus. (**I**) TSPAN8 (15 kDa) in testis, controls (males) vs. patient with Helsmoortel-Van der Aa syndrome (male), biological replicates 1-3. (**J**) Corresponding GAPDH loading-control blot for panel I. (**K**) CNN1 (30 kDa) in uterus, controls (females) vs. patient with Helsmoortel-Van der Aa syndrome (female), biological replicates 1–3. (**L**) Corresponding GAPDH loading-control blot for panel K. Molecular weight markers (kDa) are indicated on the left of each blot. The target band in each panel is boxed.

**Additional file S2. RT-PCR primers**. The table lists the SYBR Green primer sequences (5′–3′) used for RT-qPCR, indicating target gene, species, and forward/reverse primer sequence. Primers are provided for both murine and human target and reference genes.

**Additional file S3. Pyrosequencing primers**. The table lists the murine pyrosequencing primers used to quantify CpG methylation at the indicated target genes, including forward primer, reverse primer, and sequencing primer sequences (5′–3′). The biotin tag required for Sepharose bead purification is synthesised onto either the forward or reverse primer, as indicated. The sequence-to-analyse (STA) denotes the nucleotides interrogated by the Pyromark instrument. A primer score > 65 indicates proper PCR amplification.

**Additional file S4. Chronic NAP treatment does not alter body weight in WT or HET mice despite a persistent genotype-dependent difference**. (**A,B**) Body weight (grams) in male (**A**) and female (**B**) WT and HET mice following chronic DD-vehicle or NAP treatment. Under DD-vehicle conditions, HET mice of both sexes exhibited significantly lower body weight than WT littermates (male, female: p < 0.0001), and this genotype difference persisted following NAP treatment (WT + NAP vs. HET + NAP: p < 0.0001 in both sexes). Within each genotype, body weight did not differ between DD-vehicle- and NAP-treated animals (male: WT + DD vs. WT + NAP, p = 0.9940; HET + DD vs. HET + NAP, p = 0.6009; female: WT + DD vs. WT + NAP, p = 0.4648; HET + DD vs. HET + NAP, p = 0.1131), indicating that NAP treatment did not affect overall somatic growth. Each dot represents an individual animal (male: WT + DD, n = 14; HET + DD, n = 12; WT + NAP, n = 12; HET + NAP = 11; female: WT + DD, n = 11; HET + DD, n = 11; WT + NAP, n = 13; HET + NAP = 11). Horizontal lines indicate mean ± s.d. Statistical analyses were performed using two-way ANOVA followed by Tukey's multiple comparisons test.

**Additional file S5. NAP does not upregulate global or mutant *Adnp* transcript levels in HET mice**. (**A,B**) Relative *Adnp* mRNA expression normalised to *Actb*, *Gapdh*, and *Rpl13a* reference genes in male (**A**) and female (**B**) WT and HET mice following chronic DD-vehicle or NAP treatment, assessed using a global expression assay and a targeted assay specific to the CRISPR-generated frameshift mutant allele. Global *Adnp* transcript levels were reduced in HET mice relative to WT littermates under DD-vehicle conditions in both sexes (male: p = 0.0002; female: p < 0.0001), with no significant change following NAP treatment in males (p = 0.9970) and a non-significant upward trend in females (p = 0.0623). Mutant *Adnp* transcripts were detected exclusively in HET mice using the targeted assay (both sexes: p < 0.0001) and were unaffected by NAP treatment (male HET: p = 0.7591; female HET: p = 0.7818), confirming that NAP does not act by upregulating expression of the pathogenic allele. Each dot represents an individual animal (n = 7 per genotype and treatment group). Horizontal lines indicate mean ± s.d. Statistical analyses were performed using two-way ANOVA followed by Tukey's multiple comparisons test.

**Additional file S6. Sex-associated differentially methylated probes (DMPs) in the hippocampus**. Linear mixed-model ANOVA identifying CpG probes with a significant main effect of sex. Methylation β-values across all hippocampal samples were modelled with sex and genotype (*Adnp* HET versus WT) as fixed effects and no sex × genotype interaction term. Probes are filtered for a Benjamini-Hochberg (BH)-adjusted p-value < 0.05 for the sex term (n = 334 CpGs). Columns give the raw and BH-adjusted p-values for both the sex and genotype terms of the model, together with Illumina manifest probe and gene/genomic annotation.

**Additional file S7. Genotype-associated differentially methylated positions (DMPs) in the hippocampus**. Output of the same linear mixed-model ANOVA as additional file S6, here filtered for CpG probes with a nominally significant (unadjusted p < 0.05) main effect of genotype (*Adnp* HET versus WT), irrespective of sex (n = 23,873 CpGs). Columns give the raw and BH-adjusted p-values for both the sex and genotype terms of the model, together with Illumina manifest probe and gene/genomic annotation. Note that this list is filtered on the raw, not the BH-adjusted, genotype p-value.

**Additional file S8. Sex-associated methylation changes, filtered for |Δβ| ≥ 0.1**. CpG probes from the sex comparison in Additional file S6 further filtered by effect size, retaining only probes with an absolute mean β-value difference (delta-beta) between males and females of at least 0.1, independent of statistical significance (n = 993 CpGs). Columns give the raw and BH-adjusted p-values, mean β-values in males and females, delta-beta, fold change and log2 fold change, plus Illumina manifest annotation.

**Additional file S9. DMP analysis, male hippocampus only (HET vs WT), filtered for |Δβ| ≥ 0.1**. CpG probes differentially methylated between *Adnp* HET and WT male mice, tested probe-wise with a Wilcoxon rank-sum test and filtered for an absolute mean β-value difference (delta-beta) ≥ 0.1 between genotypes (n = 2,500 CpGs). Columns give the Wilcoxon p-value, BH-adjusted p-value, mean β-values in WT and HET (MUT) males, delta-beta, fold change and log2 fold change, plus Illumina manifest annotation.

**Additional file S10. DMP analysis, female hippocampus only (HET vs WT), filtered for |Δβ| ≥ 0.1**. CpG probes differentially methylated between *Adnp* HET and WT female mice, tested probe-wise with a Wilcoxon rank-sum test and filtered for an absolute mean β-value difference (delta-beta) ≥ 0.1 between genotypes (n = 1,376 CpGs). Columns give the Wilcoxon p-value, BH-adjusted p-value, mean β-values in WT and HET (MUT) males, delta-beta, fold change and log2 fold change, plus Illumina manifest annotation.

**Additional file S11. Combined-sex linear mixed-model ANOVA, genotype and NAP-treatment effects**. Linear mixed-model ANOVA testing the main effects of genotype (*Adnp* HET vs WT) and NAP treatment on hippocampal methylation with males and females analysed together and no genotype × treatment interaction term. No CpG probe reached the significance threshold for either main effect, and the file is provided as an empty results table to document that the combined-sex model detected no significant genotype- or treatment-associated DMPs. Columns give the raw and BH-adjusted p-values for the treatment and genotype (group) terms of the model, plus Illumina manifest annotation.

**Additional file S12. Combined-sex linear model ANOVA, main NAP-treatment effect**. Linear model ANOVA testing the main effect of NAP/davunetide treatment on hippocampal methylation with males and females analysed together. No CpG probe reached the significance threshold for the treatment effect, and the file is provided as an empty results table for completeness. Columns are as described for Additional file S11.

**Additional file S13. NAP treatment does not induce genome-wide changes in hippocampal DNA methylation in male and female *Adnp*-deficient mice**. (**A,B**) UMAP representations of hippocampal DNA methylation profiles in male (**A**) and female (**B**) mice treated with NAP or DD vehicle, stratified by genotype. (**C,D**) Heatmaps showing hippocampal DNA methylation profiles in male (**C**) and female (**D**) mice across WT and HET animals treated with NAP or DD vehicle. β-values are shown for CpG probes included in the analysis. No significant genotype × treatment interaction was detected at the genome-wide level, and subsequent analysis independent of genotype identified no significant treatment-associated CpG methylation changes in either sex (LMM, FDR-adjusted p > 0.01). These findings indicate that postnatal NAP treatment does not measurably alter the hippocampal DNA methylation landscape of male and female WT or HET mice.

**Additional file S14. Differentially expressed genes, male hippocampus, DD-vehicle (HET vs WT)**. Differentially expressed genes (DEGs) from bulk hippocampal RNA-seq comparing *Adnp* HET and WT male mice under DD-vehicle conditions, analysed with DESeq2. Genes are filtered for an adjusted p-value (padj, Benjamini-Hochberg) < 0.05 (n = 120 genes; 56 up- and 64 down-regulated in HET vs WT). Columns give the Ensembl gene ID and gene symbol, mean normalised counts (baseMean), log2 fold change and its standard error, Wald test p-value, and BH-adjusted p-value.

**Additional file S15. Differentially expressed genes, female hippocampus, DD-vehicle (HET vs WT)**. Differentially expressed genes (DEGs) from bulk hippocampal RNA-seq comparing *Adnp* HET and WT female mice under DD-vehicle conditions, analysed with DESeq2. Genes are filtered for an adjusted p-value (padj, Benjamini-Hochberg) < 0.05 (n = 245 genes; 163 up- and 82 down-regulated in HET vs WT). Columns give the Ensembl gene ID and gene symbol, mean normalised counts (baseMean), log2 fold change and its standard error, Wald test p-value, and BH-adjusted p-value.

**Additional file S16. Genotype × NAP-treatment interaction DEGs, male hippocampus**. Genes showing a significant genotype × treatment interaction in male hippocampal bulk RNA-seq, tested with DESeq2's likelihood-ratio test (LRT) comparing a model with a genotype × treatment interaction term to a reduced (no-interaction) model. Genes are filtered for padj < 0.05 (n = 119 genes; 56 up- and 63 down-regulated by sign of log2 fold change). Columns give the Ensembl gene ID and gene symbol, mean normalised counts (baseMean), log2 fold change and its standard error, LRT p-value, and BH-adjusted p-value, with the addition of the test statistic (stat) from the interaction term.

**Additional Figure S17. Genotype-dependent transcriptional response to NAP treatment in the hippocampus of male *Adnp*-deficient mice**. (**A**) Heatmap showing representative transcripts with a significant genotype × treatment interaction in male mice across WT and HET littermates treated with DD vehicle or NAP. Samples were hierarchically clustered according to their expression profiles. Genotype and treatment are indicated above the heatmap. (**B**) Functional enrichment analysis of genes showing a significant genotype-dependent response to NAP treatment in male mice (LRT, FDR < 0.05). Enriched pathways are shown according to the direction of the response, with suppressed pathways including cell-cycle, mitotic regulation, and mRNA-splicing processes, and activated pathways including neurotransmitter receptor and synaptic signalling, AMPA receptor trafficking, and RHO GTPase cycling. Dot size represents the number of genes contributing to each pathway and dot size and colour indicate gene count and adjusted p-value, respectively. (**C**) RT-qPCR validation of *Map2* expression in male WT and HET mice treated with DD vehicle or NAP. *Map2* expression did not differ between genotypes under DD vehicle treatment but was significantly higher in HET than in WT mice following NAP treatment. NAP treatment also significantly increased *Map2* expression in HET mice. Data are presented as mean ± s.d. and were analysed using two-way ANOVA with Šídák's multiple comparisons test (male WT + DD, n = 5; male HET + DD, n = 4; male WT + NAP, n = 5; male HET + NAP, n = 4).

**Additional file S18. Genotype × NAP-treatment interaction DEGs, female hippocampus**. Genes showing a significant genotype × treatment interaction in female hippocampal bulk RNA-seq, tested with DESeq2's likelihood-ratio (LRT) test comparing a model with a genotype × treatment interaction term to a reduced (no-interaction) model. Genes are filtered for padj < 0.05 (n = 4 genes; 2 up- and 2 down-regulated by sign of log2 fold change). Columns give the Ensembl gene ID and gene symbol, mean normalised counts (baseMean), log2 fold change and its standard error, LRT p-value, and BH-adjusted p-value, with the addition of the test statistic (stat) from the interaction term.

**Additional file S19. Hippocampal expression of hormone-associated genes in female and male mice**. Expression of *Esr1*, *Pgr*, and *Bdnf* was assessed by bulk RNA sequencing in hippocampal samples from female and male mice treated with DD vehicle or NAP. Female mice were compared with male mice as a non-cycling reference to assess whether the female cohort showed a pronounced high-oestrogen transcriptional state at tissue collection. No differences in expression of *Esr1*, *Pgr*, and *Bdnf* were observed in female mice at either genotype or treatment condition compared with the corresponding male group. Data are presented as normalised read counts (male: WT + DD, n = 5, HET + DD, n = 4; WT + NAP, n = 5, HET + NAP, n = 4; female: WT + DD, n = 6, HET + DD, n = 5; WT + NAP, n = 5, HET + NAP, n = 5). Statistical analysis was performed using the likelihood-ratio (LRT) test with FDR correction (FDR < 0.05). FDR-adjusted p-values are shown.

**Additional file S20. ChEA3 transcription factor enrichment analysis of 4CG hippocampal DEGs**. ChEA3 (ChIP-X Enrichment Analysis version 3) transcription factor enrichment results generated from the differentially expressed genes identified in the Four Core Genotypes (4CG) hippocampal RNA-seq analysis. Transcription factors are ranked by their integrated (mean) ChEA3 rank across the underlying reference libraries (ARCHS4 Coexpression, ENCODE ChIP-seq, ReMap ChIP-seq, Literature ChIP-seq, Enrichr Queries and GTEx Coexpression), from most (rank 1) to least enriched (rank 1631); ADNP itself ranks 219/1631. Columns give the query name, integrated rank, TF symbol, ChEA3 score, the contributing libraries with their individual (per-library) rank, and the overlapping query genes supporting each TF's enrichment.

**Additional file S21. Molecular characterisation of ADNP and representative reproductive-tissue transcriptional and protein changes in human Helsmoortel-Van der Aa syndrome**. (**A,B**) Global transcript analysis of *ADNP* and representative genes identified from the mouse transcriptomic analysis in human testicular (**A**) and uterine (**B**) tissue from control subjects and individuals with Helsmoortel-Van der Aa syndrome. *ADNP* expression was assessed for total transcript abundance. Representative mouse-associated transcripts included *ADH1C* and *TSPAN8* in testis and *CNN1* and *TGFBR3L* in uterus. *TSPAN8* and *CNN1* showed changes in the same direction as observed in the corresponding mouse tissues, whereas *ADH1C* and *TGFBR3L* showed changes in the opposite direction. Relative mRNA expression was normalised to *TBP*, *RPL13A*, and *RPLP0*. Statistical analysis was performed using Welch's t-test. Three independent control individuals were used for RNA extraction, whereas three independent biopsy samples were obtained from one male and one female individual with Helsmoortel-Van der Aa syndrome. (**C,D**) Representative immunohistochemical staining and quantification of TSPAN8 in human testicular tissue (**C**) and CNN1 in human uterine tissue (**D**) from control subjects and individuals with Helsmoortel-Van der Aa syndrome. Statistical analysis was performed using Welch's t-test. (**E**) Representative western blot analysis and quantification of TSPAN8 in testicular tissue and CNN1 in uterine tissue from control subjects and individuals with Helsmoortel-Van der Aa syndrome. Data are presented as mean ± s.d. Statistical analysis was performed using Welch's t-test.
