## Supplementary figures and images for "The Female Side of Autism: Sexual Dichotomies impacting the Helsmoortel-Van der Aa syndrome pathology"

### S1

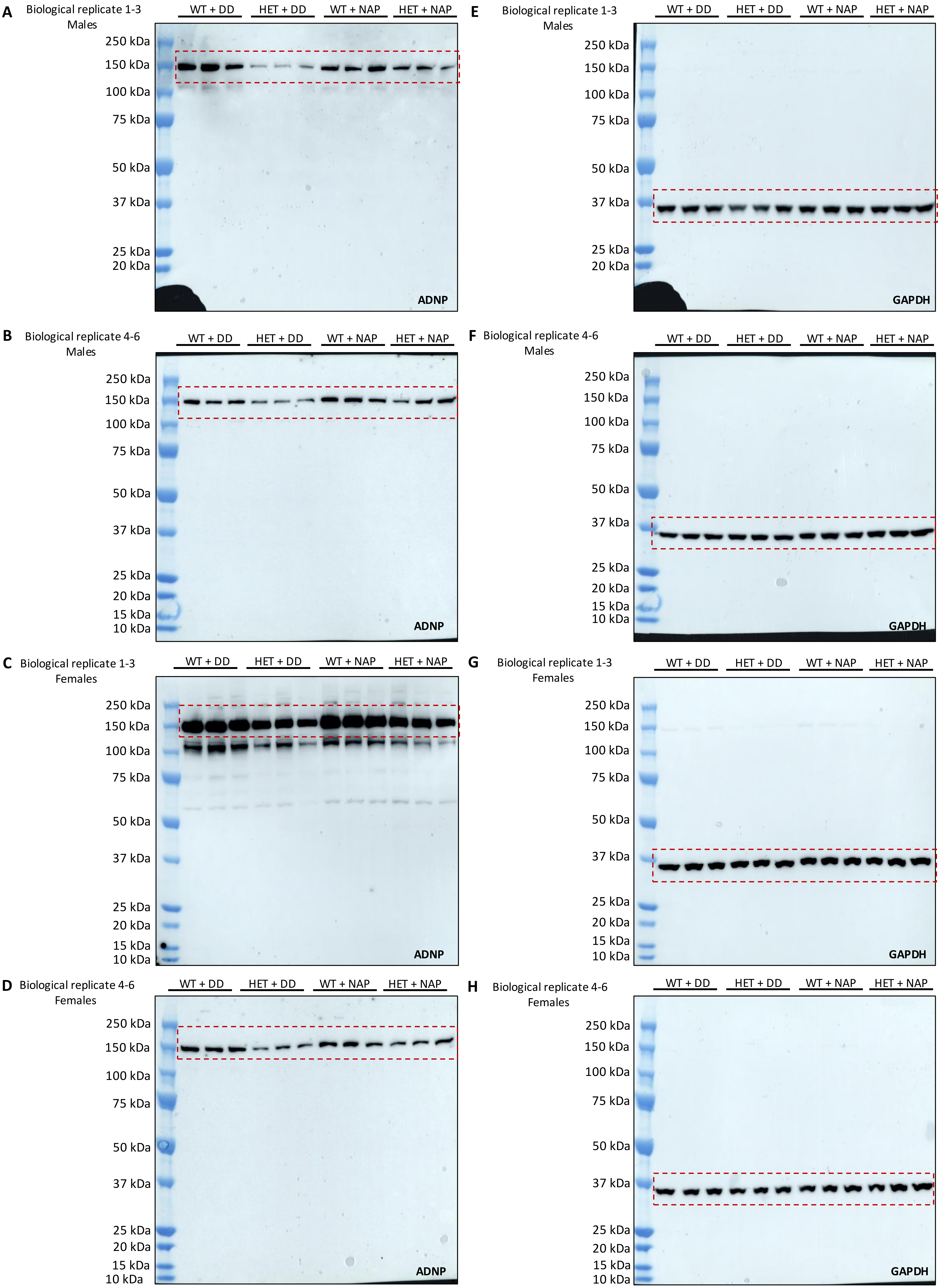

**I** Testis replicate 1-3

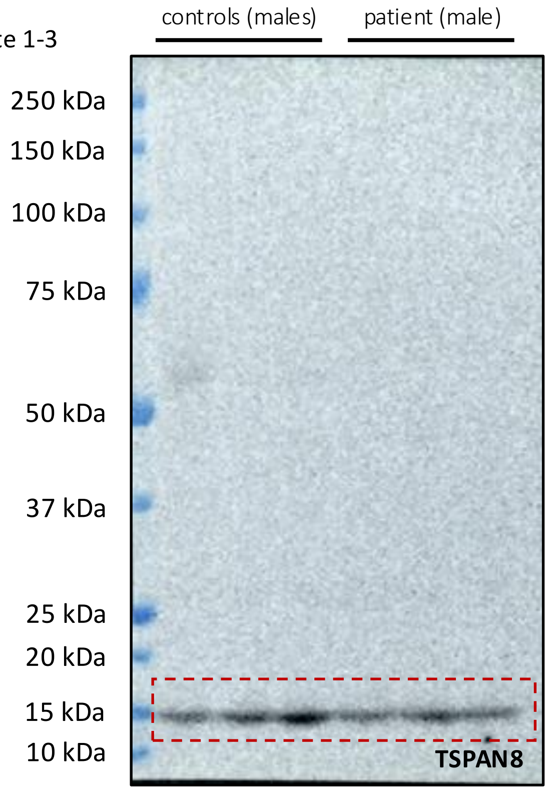


**K** Uterus replicate 1-3

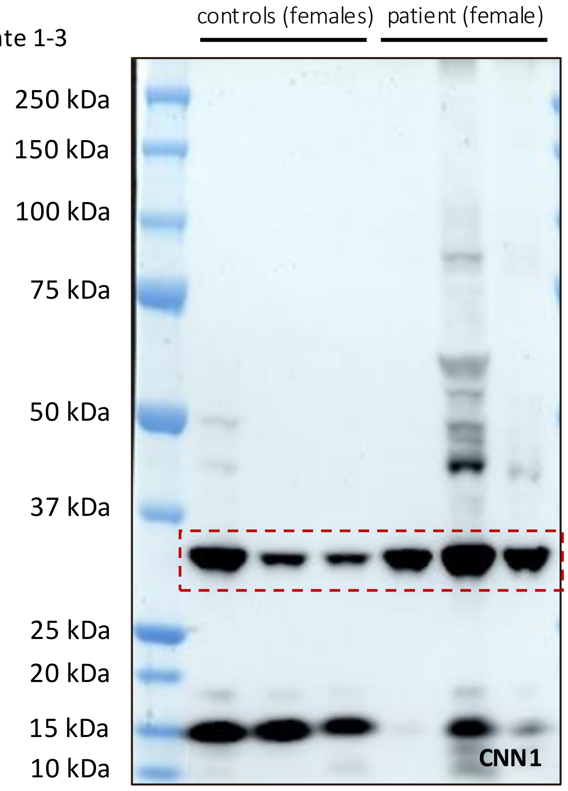


**J** Testis replicate 1-3

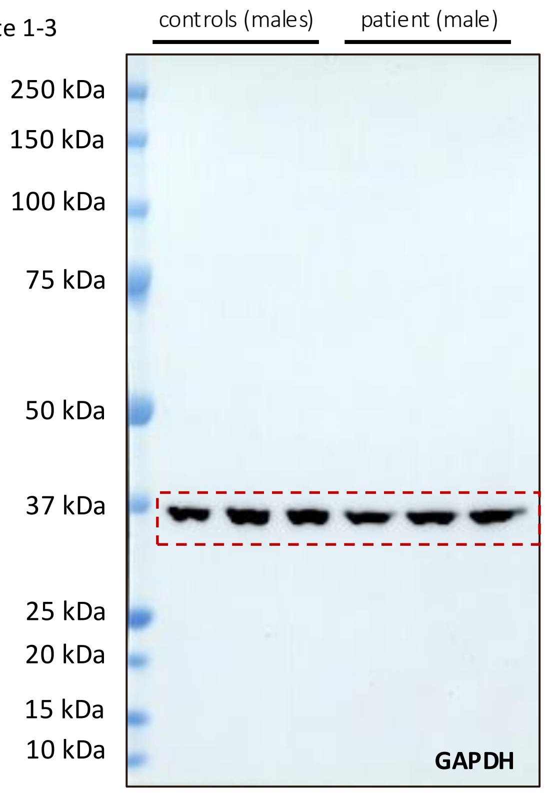


**L** Uterus replicate 1-3

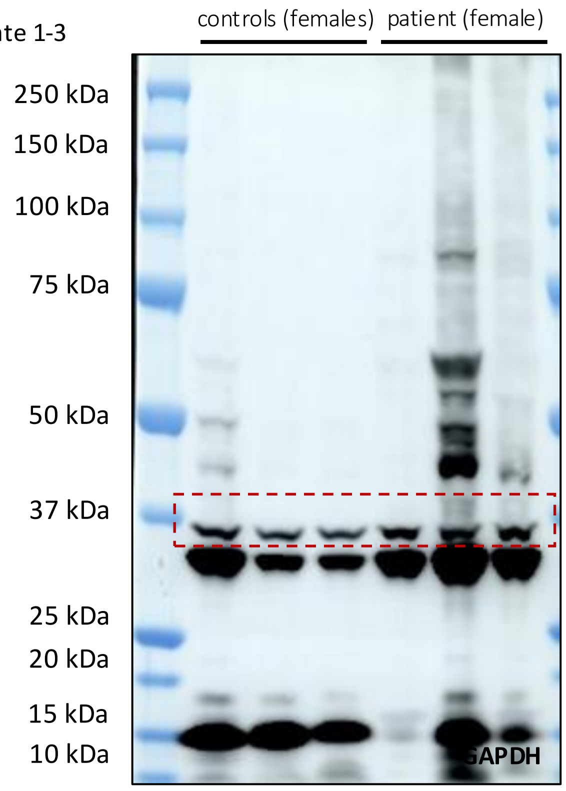

### S4

**Additional file S4, related to figure 3**

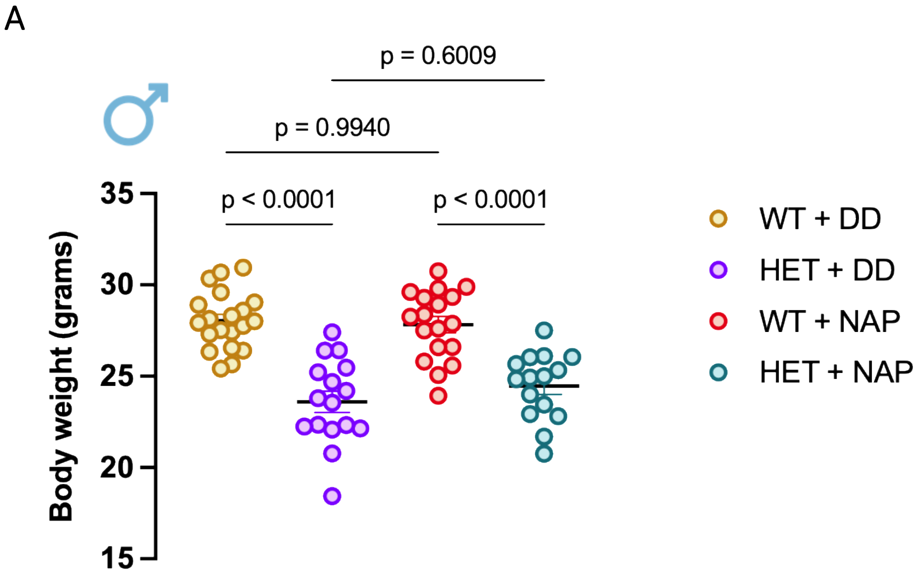


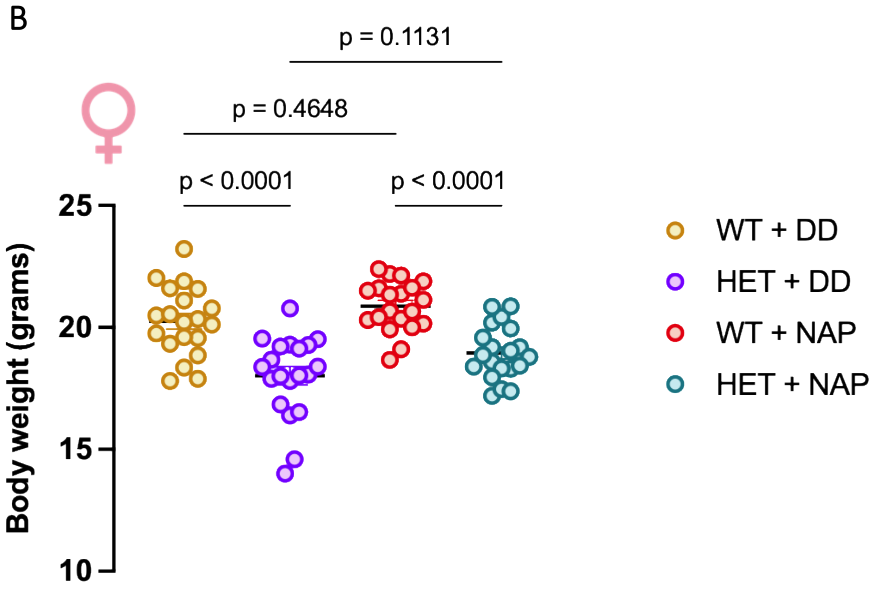

### S5

#### Additional file S5, related to figure 3

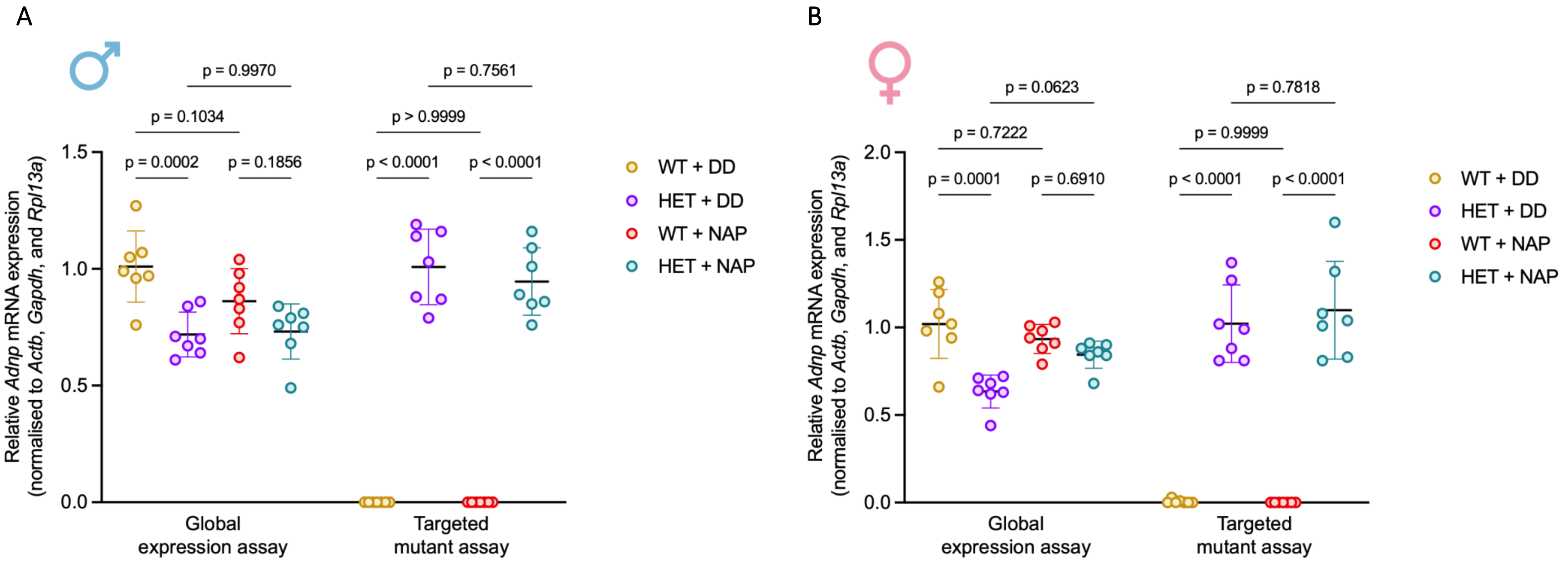

### S13

**Additional file S13, related to main figure 5**

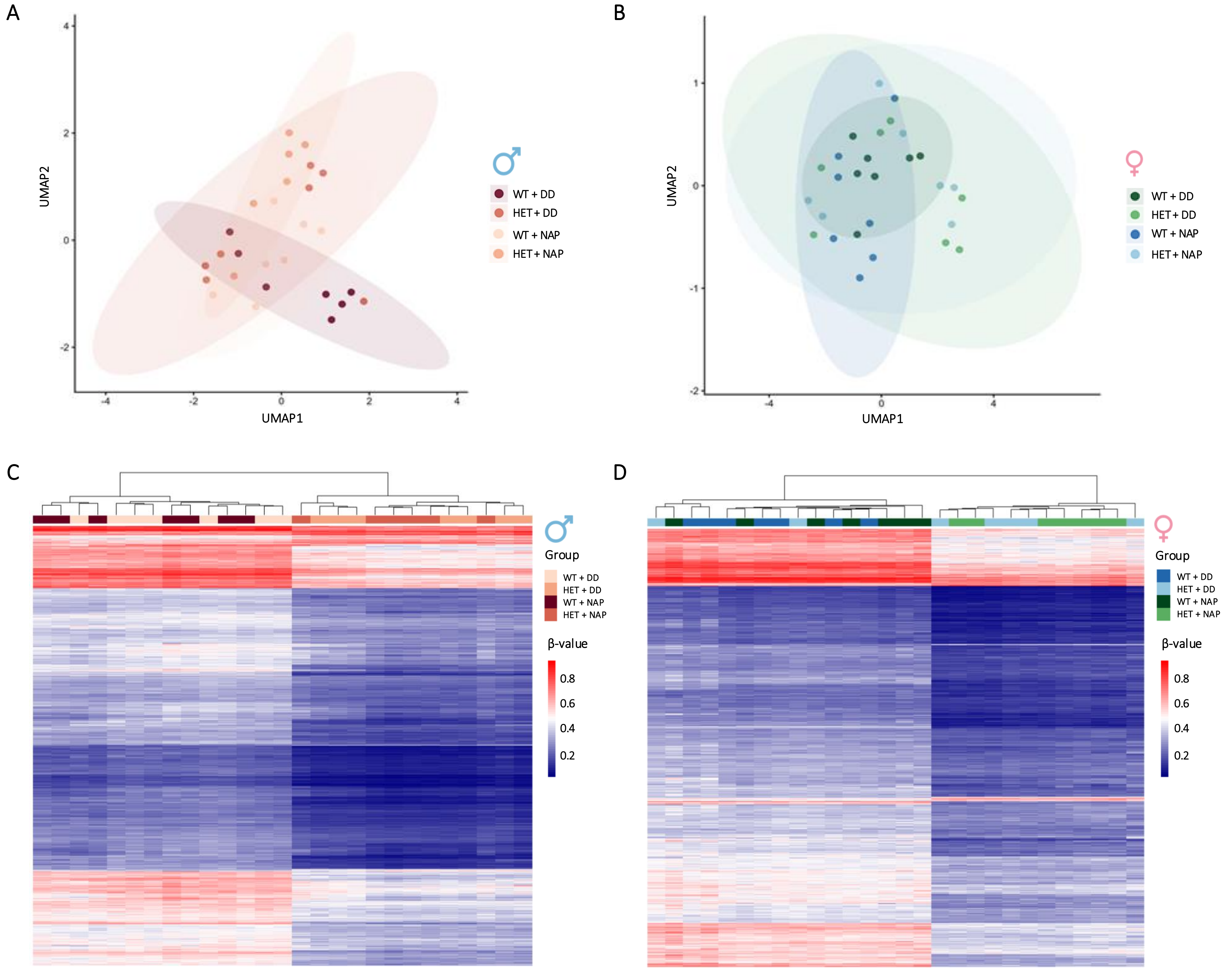

### S19

**Additional file S19, related to main figure 6**

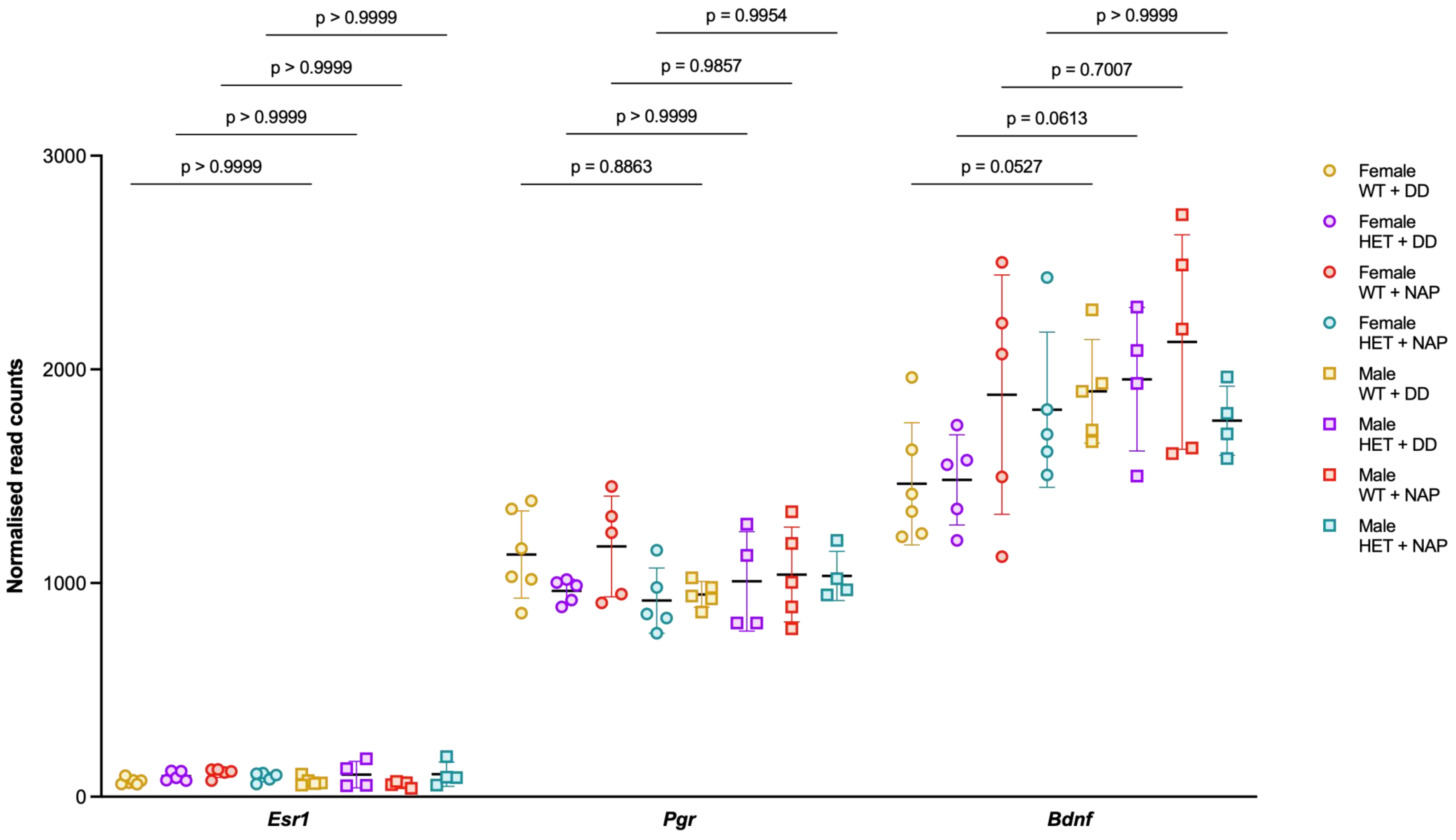

### S21

#### Additional file S21, related to main figure 7

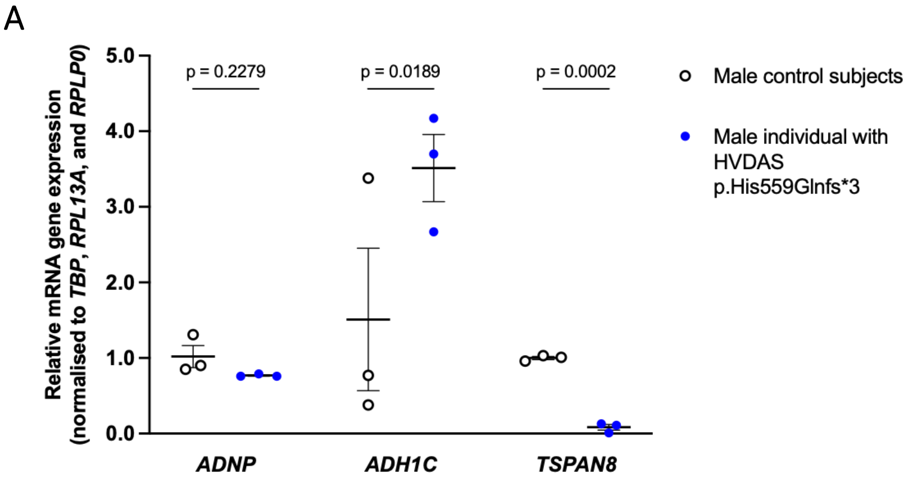


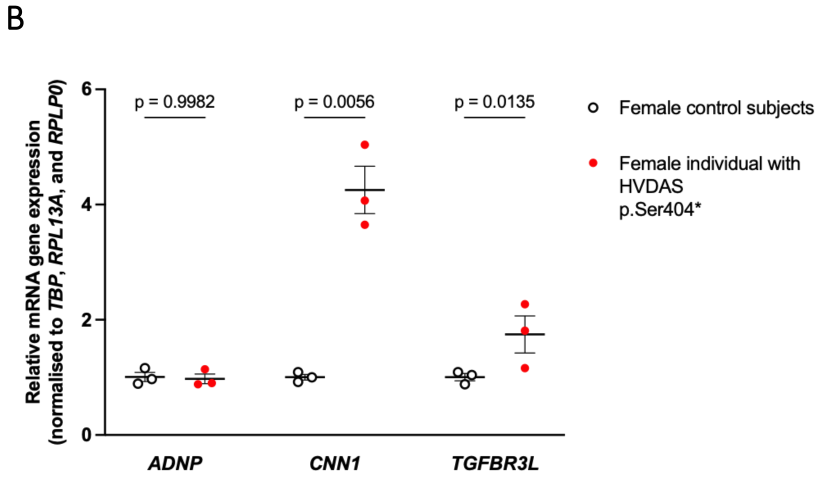


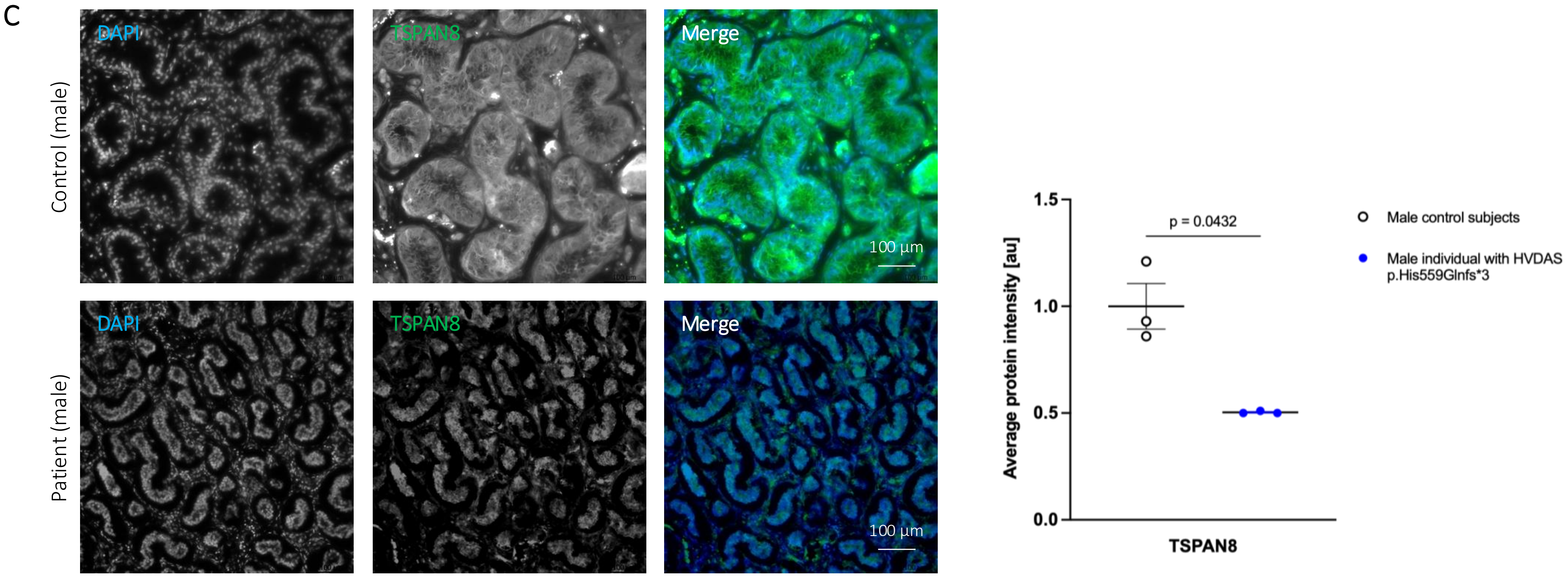


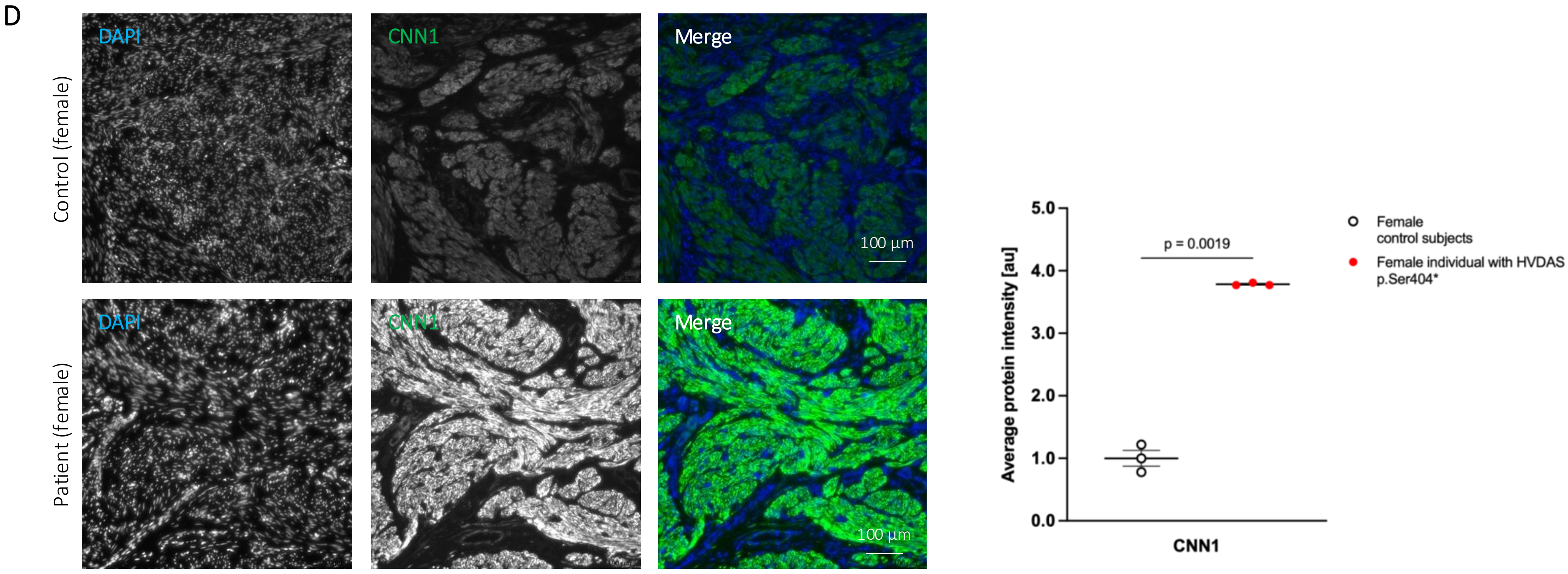


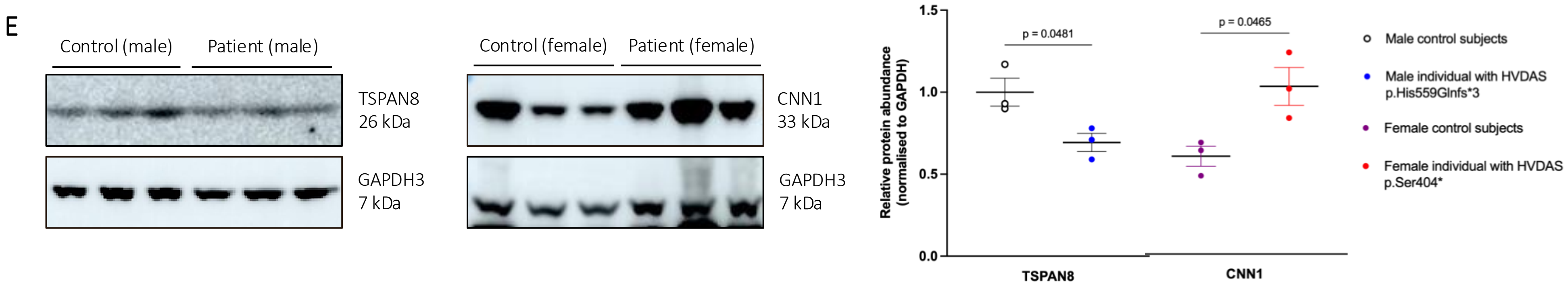
