## Supplementary material for "The Female Side of Autism: Sexual Dichotomies impacting the Helsmoortel-Van der Aa syndrome pathology": S2

**Additional file S2. RT-PCR primers**. The table lists the SYBR Green primer sequences (5′–3′) used for RT-qPCR, indicating target gene, species, and forward/reverse primer sequence. Primers are provided for both murine and human target and reference genes.

| Target gene | Species | Forward primer | Reverse Primer |
| --- | --- | --- | --- |
| *Nlgn1* | Mus musculus | ACAGGAGAACATCGTTTCCAGCCT | ATACAGGAGCAAACTGAGTGGCGT |
| *Rmi1* | Mus musculus | CACAACAGACCCTCAGCAGT | CTTGCAATTCCAGCACCACC |
| *Map2* | Mus musculus | GAAGTTCAGGCCCACTCTCC | ATTCTTCAGGTCCGGCAGTG |
| *Eif4b* | Mus musculus | GGGTGGTGAATAGGGGTGTG | ACAGCTGTCTTCGGATGCAA |
| *Global Adnp expression assay (exon 5_no deletion)* | Mus musculus | AACGGCAATACGGACAAGAC | GCGTTCACTTCGAAAAGGAG |
| *Targeted Adnp wild-type assay* | Mus musculus | CGTGATTTTCAAACAGCCACTC | GCCTGGTGTGCTGCTAGGTT |
| *Targeted Adnp mutant assay* | Mus musculus | TTCCCAAGGTTATGCTTTTTGT | GCCTGGTGTGCATGAAAG |
| *Col6a4* | Mus musculus | AAGAGGATTTTCAGGAGAGAAGGG | AGATTATCAATTCCAGGATCCCCC |
| *Card10* | Mus musculus | CACCATCCTTGTGACACTGG | GGAATCACCCACAAGCCTAA |
| *Homer3* | Mus musculus | TTCCAGATCGACCCCACTAC | GGGAGTGACAGTGCTGTTGA |
| *Satb2* | Mus musculus | GCAGTTAAGCCAGAGCCAAC | GCAGGTTGAGGAAGTTCTGC |
| *Adh1* | Mus musculus | CCGGTTCACTAGAGGAGGGA | AGCTCCATCGATTTTGGCCA |
| *Tspan8* | Mus musculus | TCTCTGCAGGCAAAATCCGT | CCAAGCCACAGCACTTGAAC |
| *Cnn1* | Mus musculus | GTGGATTGAGGGGGTGACAG | ACCTGGCTCAAAGATCTGCC |
| *Tgfbr3l* | Mus musculus | GTCAGCTTCTCACCACCTCC | GCCGCTGAGAGAAGCTCTAG |
| *Gapdh* | Mus musculus | GGCATTGCTCTCAATGACAA | CCCTGTTGCTGTAGCCGTAT |
| *β-Actin* | Mus musculus | CAACGAGCGGTTCCGATG | GCCACAGGATTCCATACCCA |
| *Rpl13a* | Mus musculus | GCAGGCATGAGGCAAACAGTC | CACTCTGGAGGAGAAACGGAAGG |
| *B2m* | Mus musculus | ATGCACGCAGAAAGAAATAGCAA | AGCTATCTAGGATATTTCCAATTTTTTGAA |
| *Global Adnp expression assay* | Homo sapiens | CACCTTGCATGGTAGCCTTT | TGAGGTTGACCAAGACGATG |
| *ADH1C* | Homo sapiens | GATGGTGGCTGCAGGAATCT | GGCACCCAACTCTTTAGCCT |
| *TSPAN8* | Homo sapiens | TGCCTGGAGATAGCCTTTGC | CTGTGGCGCTCAAAAGCTTT |
| *CNN1* | Homo sapiens | TTGAGGCCAACGACCTGTTT | CTGGGTACTCGGGAGTCAGA |
| *TGFBR3L* | Homo sapiens | GACACCTCTGTCGCCTTCC | TGGGTGTATCTCCGGACCAT |
| *TBP* | Homo sapiens | GCAAGGGTTTCTGGTTTGCC | CAAGCCCTGAGCGTAAGGTG |
| *RPL13A* | Homo sapiens | CCTGGAGGAGAAGAGGAAAGAGA | TTGAGGACCTCTGTGTATTTGTCAA |
| *RPLP0* | Homo sapiens | CCTCGTGGAAGTGACATCGT | CTGTCTTCCCTGGGCATCAC |
