## Supplementary material for "The Female Side of Autism: Sexual Dichotomies impacting the Helsmoortel-Van der Aa syndrome pathology": S3

**Additional file S3. Pyrosequencing primers**. The table lists the murine pyrosequencing primers used to quantify CpG methylation at the indicated target genes, including forward primer, reverse primer, and sequencing primer sequences (5′–3′). The biotin tag required for Sepharose bead purification is synthesised onto either the forward or reverse primer, as indicated. The sequence-to-analyze (STA) denotes the nucleotides interrogated by the Pyromark instrument. A primer score > 65 indicates proper PCR amplification.

| Gene | Forward primer | Reverse primer | Sequencing primer | Biotin tag | Sequence-to-analyze | Score |
| --- | --- | --- | --- | --- | --- | --- |
| *Gsk3b*  *(hypo in both males and female mice)* | TTTTGGTTTATGTGTTATGTATTTGTGAGT | CTACATTTTCCTATTCCTCAAATAATCT | GTATTTGTGAGTTAGATGTT | Rev | TTYGTGTTTT GGTAG | 80 |
| *Apoa1*  *(hypo*  *in male mice)* | GTATTGATTTGGATAGTGGAGTATTAATTG | CCTACCCACACACATATATAAACC | AGGAGTTATGGAGTTAGAGTTAT | Rev | YGAAGGTAGG TAGTAGTATT TATTTT | 87 |
| *Foxp2*  *(hypo*  *in female mice)* | AGAGAGAGGTTTATTTTAAGAAGTGTA | AAACAAACTCATTTTCCTATCCTAATAC | AAAGTTGGATAGTTATTGTT | Rev | TAGTTATTTY GTAGTAATTT TATGGAGTAT TAGGA | 89 |
