## Supplementary material for "The Female Side of Autism: Sexual Dichotomies impacting the Helsmoortel-Van der Aa syndrome pathology": S17

### Additional file S17, related to main figure 6

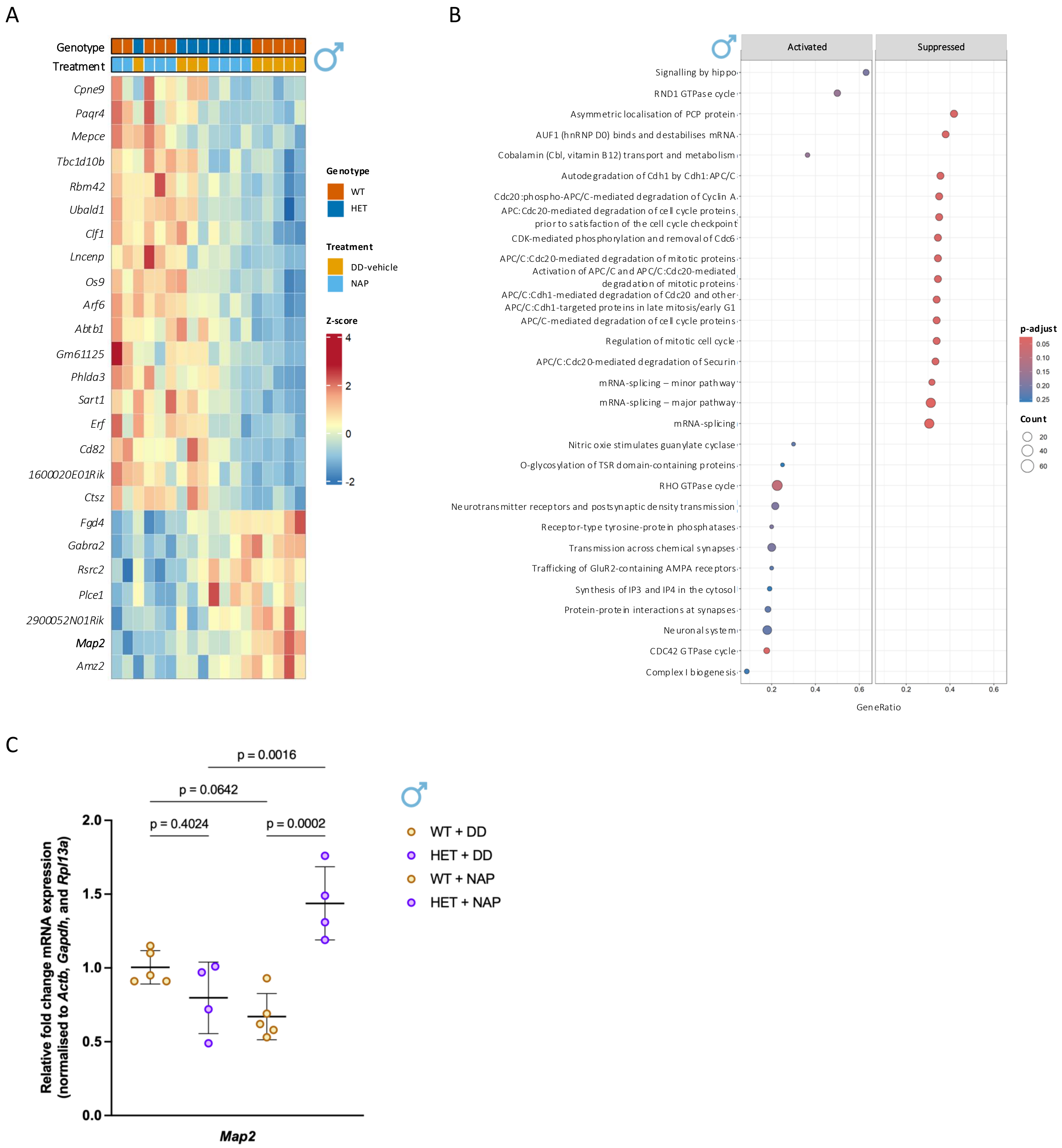


**Additional Figure S17. Genotype-dependent transcriptional response to NAP treatment in the hippocampus of male *Adnp*-deficient mice.** (**A**) Heatmap showing representative transcripts with a significant genotype × treatment interaction in male mice across WT and HET littermates treated with DD vehicle or NAP. Samples were hierarchically clustered according to their expression profiles. Genotype and treatment are indicated above the heatmap. (**B**) Functional enrichment analysis of genes showing a significant genotype-dependent response to NAP treatment in male mice (FDR < 0.05). Enriched pathways are shown according to the direction of the response, with suppressed pathways including cell-cycle, mitotic regulation, and mRNA-splicing processes, and activated pathways including neurotransmitter receptor and synaptic signalling, AMPA receptor trafficking, and RHO GTPase cycling. Dot size represents the number of genes contributing to each pathway and dot size and colour indicate gene count and adjusted p-value, respectively. (**C**) RT-qPCR validation of *Map2* expression in male WT and HET mice treated with DD vehicle or NAP. *Map2* expression did not differ between genotypes under DD vehicle treatment but was significantly higher in HET than in WT mice following NAP treatment. NAP treatment also significantly increased *Map2* expression in HET mice. Data are presented as mean ± s.d. and were analysed using two-way ANOVA with Šídák's multiple comparisons test (male WT + DD, n = 5; male HET + DD, n = 4; male WT + NAP, n = 5; male HET + NAP, n = 4).
